# *Fancm* regulates meiotic double-strand break repair pathway choice in mammals

**DOI:** 10.1101/2022.06.16.496499

**Authors:** Vanessa Tsui, Ruqian Lyu, Stevan Novakovic, Jessica M. Stringer, Jessica E. M. Dunleavy, Elissah Granger, Tim Semple, Anna Leichter, Luciano G. Martelotto, D. Jo Merriner, Ruijie Liu, Lucy McNeill, Nadeen Zerafa, Eva Hoffmann, Moira K. O’Bryan, Karla Hutt, Andrew J. Deans, Jörg Heierhorst, Davis J. McCarthy, Wayne Crismani

**Author notes:** Correspondence to Wayne Crismani, Davis McCarthy. The authors wish it to be known that, in their opinion, the first two authors should be regarded as joint first authors.

## Abstract

Meiotic crossovers are required for accurate chromosome segregation and to produce new allelic combinations. Meiotic crossover numbers are tightly regulated within a narrow range, despite an excess of initiating DNA double-strand breaks. Here, we describe the tumour suppressor FANCM as a meiotic anti-crossover factor in mammals. Crossover analyses with single-gamete and pedigree datasets both reveal a genome-wide increase in crossover frequencies in *Fancm*-deficient mice. Gametogenesis is heavily perturbed in *Fancm* loss of function mice, which is consistent with the reproductive defects reported in humans with biallelic *FANCM* mutations. A portion of the gametogenesis defects can be attributed to the cGAS-STING pathway. Despite the gametogenesis phenotypes in *Fancm* mutants both sexes were capable of producing offspring. We propose that the anti-crossover function and role in gametogenesis of *Fancm* are separable and will inform diagnostic pathways for human genomic instability disorders.

## Introduction

Fanconi anaemia (FA) is a rare genetic condition which predisposes to health issues such as cancers, progressive bone marrow failure, birth defects and reduced fertility [1, 2]. The genes which are mutated in individuals with FA act in a common genetic and biochemical pathway. The FA pathway is a DNA repair mechanism best known for its role in the repair of DNA inter-strand crosslinks in somatic cells. There are 22 “FANC” genes (FANCA-FANCW) [3]. FANCM is a DNA translocase, which can remodel branched DNA structures and provides a physical connection between multiple genomic stability-related proteins complexes. FANCM orthologues are widely conserved across eukaryotes and prokaryotes. However, vertebrates tend to have an additional ERCC4 domain at the carboxyl terminus compared to other taxa [4]. The significance of the ERCC4 in vertebrates is not clear as it appears to lack catalytic activity.

In humans and mice, FANCM is required for normal fertility and gametogenesis [5–12]. Reports of male patients with biallelic mutations in *FANCM* revealed sterility characterised by non-obstructive azoospermia and Sertoli cell-only seminiferous epithelium [8, 9, 13]. Females with a homozygous pathogenic *FANCM* mutation experienced premature ovarian insufficiency and therefore early menopause [10]. It was previously reported that a reduction in the number of primordial germ cells occurs prior to birth in *Fancm* mutant mice of both sexes [6]. Similar losses of primordial germ cells have been observed in other ’*Fanc*’ mouse models [14–16]. In *Fancm*-deficient mice, at the molecular level a significant portion of this reduction was attributed to the ATM-p53-p21 axis, most notably in the testis. Further, female *Fancm* mutants were born at subMendelian ratios [5–7] and this was attributed to embryonic death induced by genomic instability activating inflammatory pathways [7]. Males pups, on the other hand, were born at the normal frequencies [7].

A key process within germ cell development is meiosis. At meiosis, crossovers (COs)—large reciprocal exchanges between homologous chromosomes—are necessary for balanced chromosome segregation at the first division. Strict mechanisms exist to ensure that each pair of homologous chromosomes forms at least one “obligate” crossover [17]. An absence of crossovers between homologues can lead to aneuploid karyotypes and conditions such as trisomy 21 [18, 19]. The obligate crossover is linked to a phenomenon known as crossover interference, which results in relatively even spacing of multiple crossovers on the same chromosome [20–23]. Additional layers of control of crossover distribution are determined by PRDM9 through the direction of the initiating DNA double-strand breaks (DSBs) [24–26] as well as separate mechanisms that influence megabase-scale crossover patterning [27–30].

Meiotic crossovers are formed via the highly regulated and well conserved meiotic DSB repair pathway [17]. Programmed meiotic DNA DSBs are generated by SPO11 [31–33]. The DSBs are resected to generate 3*^1^* single-strand overhangs [34–37]. The recombinases RAD51 and DMC1 are recruited to these single-stranded tails and form nucleoprotein filaments, which promote repair that uses the homologue as a template to fill in missing genetic information. The single-stranded nucleoprotein filament displaces one strand of the homologue forming a displacement loop (D-loop) [35, 38, 39]. Second-end capture can progress the D-loop towards “joint molecules” such as a double Holliday junction, which is dependent on a group of proteins collectively referred to as the ZMMs (for reviews see [40–42]). The ZMMs are required for most crossovers in animals, budding yeast and plants, and these crossovers are sensitive to interference and often referred to as “class I” crossovers [41, 43–46]. The number of class I crossovers is positively correlated with synaptonemal complex length [47–50]. An alternative “class II” crossover pathway produces a smaller proportion of crossovers that are not sensitive to interference [46, 51–57].

SPO11-dependent meiotic DSBs occur in excess to the number of crossovers in diverse species and therefore conserved mechanisms likely exist that cap the number of crossovers per meiosis [20, 21]. When programmed meiotic DSBs are repaired with a homologous template, joint molecules are formed as intermediates prior to a crossover or non-crossover outcome [42, 43, 58–63], and therefore joint molecules present a key step at which crossover rates can be altered. FANCM family members have previously been shown to dissociate joint molecules with high efficiency via helicase activity in yeast [64, 65] and translocase activity in mammals [66]. Additionally, FANCM, along with co-factors MHF1 and MHF2 has been found to limit the number of meiotic crossovers in *Arabidopsis*, fission yeast, Drosophila, canola, rice and pea [67–74]. This crossover-limiting function of FANCM may occur through the ability to dissociate joint molecules during meiotic DSB repair, but is difficult to demonstrate directly. However, any potential role of *Fancm* in meiotic crossover regulation in mammals remains unexplored.

Here we reveal an anti-crossover role during meiosis of *Fancm* in mammals using traditional pedigree-based techniques and single-sperm sequencing methods. We also build on previous findings on the role of *Fancm* in fertility and gametogenesis. We reveal a role for the cGAS-STING innate immune response in driving the loss of germ cells in males, which indicates that the cGAS-STING pathway may be a target for ameliorating the impaired gametogenesis that occurs in patients with genomic instability.

## Results

### Generation of *Fancm* mutant mice

To investigate a potential function for *Fancm* in crossover control in mammals, we generated two mouse models. *Fancm* knockout mice were generated independently – with two separate targeting events – in different genetic backgrounds, FVB/N and C57BL/6J, herein referred to as FVB.*Fancm^Δ^*^2/^*^Δ^*^2^ and B6.*Fancm^Δ^*^2/^*^Δ^*^2^, respectively (Supplementary Fig. 1). Two sgRNAs were generated to target the second exon of *FANCM*, similar to a previous approach [5]. Targeted resequencing of the exon 2 borders confirmed the deletion (Supplementary Fig. 1). The *Fancm* heterozygous founders were backcrossed three times to their respective strains of FVB/N or C57BL/6J to increase the probability of eliminating any possible off-target mutations that may have been introduced by Cas9. *Fancm^Δ^*^2/^*^Δ^*^2^ mice were viable in both strains. Null mice of both strains were outwardly normal except for a slight reduction in the body weight of female B6.*Fancm^Δ^*^2/^*^Δ^*^2^ mice (p < 0.0001, Supplementary Fig. 2), and sub-fertility. Complete blood counts were similar to the wild-type controls (Supplementary Fig. 2). Cancers were not detected in any mice in the study, however mice used in the study were typically young adults. The loss of *Fancm* resulted in phenotypes consistent with what has been observed previously [5, 6, 8]. We observed a dose-dependent increase in chromosome fragility in lymphocytes following exposure to the cross-linking drug mitomycin C (Supplementary Fig. 3a, b). We also observed an increase in micronucleated red blood cells in basal conditions similar to previous findings [6] (Supplementary Fig. 3c). Finally, we observed a loss of FANCD2 mono-ubiquitination in splenic B cells in basal conditions, under replication stress (hydroxyurea) and when treated with a DNA crosslinker (mitomycin C), consistent with loss of function of *Fancm* and the loss of the catalytic function of the E3 ligase complex that it anchors to somatic DNA damage [5, 75] (Supplementary Fig. 3d).

**Fig. 1:**
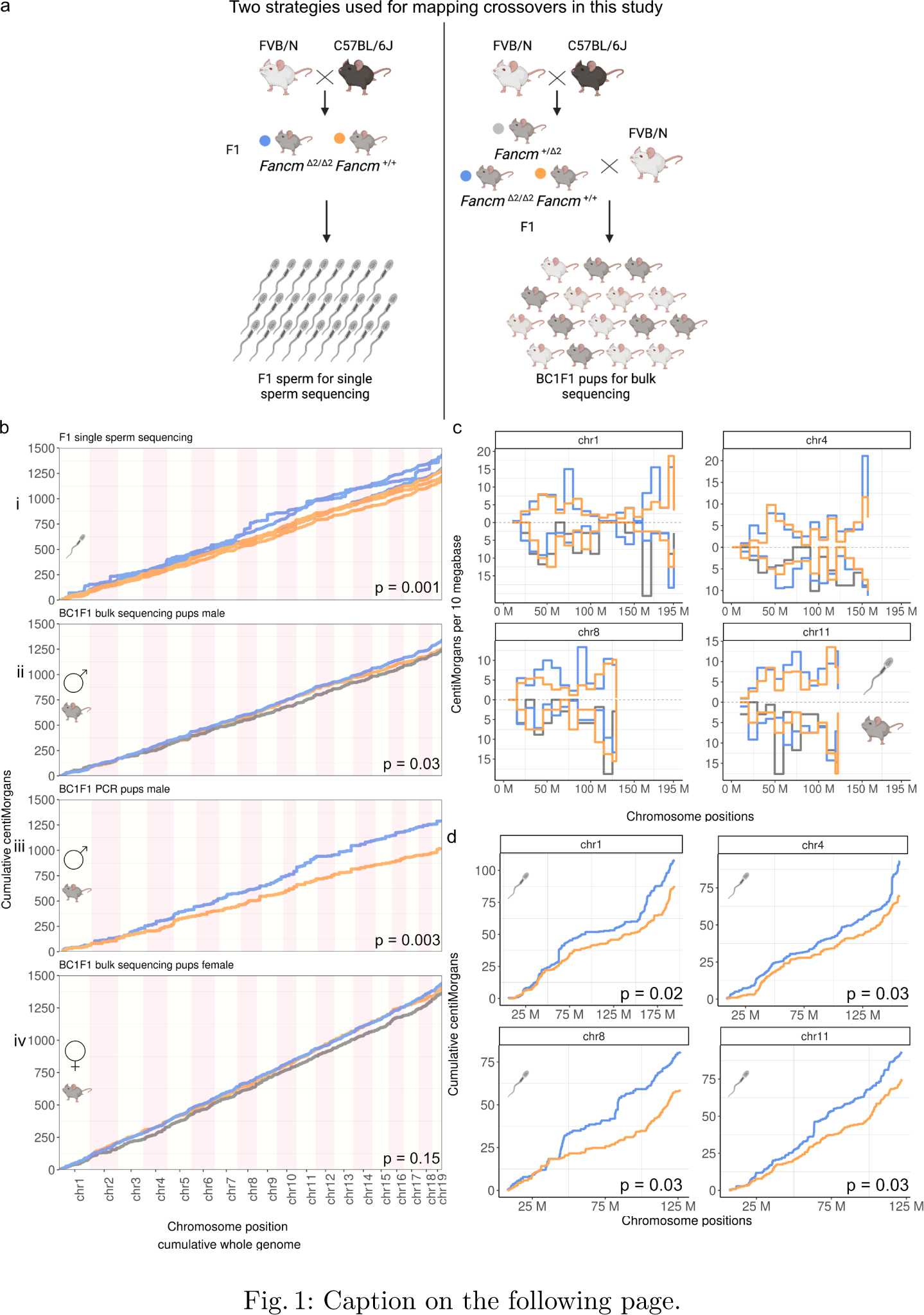
*Fancm* limits meiotic crossovers as measured using different generations and technologies. a) Crossovers were assayed using high-throughput sequencing of individual sperm from F1 mice, bulk whole-genome DNA sequencing of BC1F1 animals and PCR of selected SNPs. b) Cumulative centiMorgans calculated using data from three different assays; i) F1 single sperm sequencing (n = 3 animals per genotype, n = 218 sperm from *Fancm^Δ^*^2/^*^Δ^*^2^, 190 sperm from *Fancm*^+/+^); ii) high-throughput sequencing with BC1F1 pups (n = 98 mice from *Fancm^Δ^*^2/^*^Δ^*^2^, 34 mice from *Fancm*^+/^*^Δ^*^2^, 40 mice from *Fancm*^+/+^); and iii) PCR-based markers with BC1F1 pups (n = 31 *Fancm^Δ^*^2/^*^Δ^*^2^, 25 *Fancm*^+/+^ independent animals per genotype). iv) 200 BC1F1 pups sequenced to derive female genetic distances (n = 100 *Fancm^Δ^*^2/^*^Δ^*^2^, 50 *Fancm*^+/^*^Δ^*^2^, 50 *Fancm*^+/+^ independent animals per genotype). Observed crossover fractions were used as input and converted into genetic distances in centiMorgans using the Kosambi mapping function and presented as cumulative centiMorgans across the genome. All three approaches demonstrate a significant increase in total genetic map lengths in male *Fancm^Δ^*^2/^*^Δ^*^2^ mutants compared to that measured in *Fancm*^+/+^ littermates. Note: for PCR genotyping chromosome 10 is not presented as full length due to lower marker coverage. c,d) Four selected chromosomes 1, 4, 8 and 11 which drove a significant portion of the increase in map length in the mutants are plotted. c) Genetic distances were plotted in 10 Mb chromosome bins for four selected chromosomes for two assays (F1 single sperm sequencing (top) and bulk DNA sequencing of BC1F1 pups (bottom)). No significant differences were observed in genotype groups in each bin. d) Cumulative genetic distances per chromosome from F1 single sperm sequencing data with permutation p values printed for each chromosome after multiple testing correction.

### *Fancm* mutant mice are sub-fertile

Human studies have reported that loss of *FANCM* is associated with infertility in males and premature menopause in females [8, 10]. Therefore, in order to extend our understanding of the origins of this pathology using mouse models, we set up breeding pairs with either a female or male *Fancm^Δ^*^2/^*^Δ^*^2^ mouse with a *Fancm*^+/+^ or *Fancm*^+/^*^Δ^*^2^ partner. Both male and female *Fancm^Δ^*^2/^*^Δ^*^2^ produced pups that appeared outwardly normal. Average litter sizes were significantly reduced in male and female B6.*Fancm^Δ^*^2/^*^Δ^*^2^ mice (p = 0.004 and p = 0.022, respectively) compared to B6.*Fancm*^+/^*^Δ^*^2^ controls (Supplementary Fig. 4a). The average litter sizes were not different between *Fancm* loss of function genotypes for FVB/N and F1 strains. There was no difference observed in the average time between litters for all strains (Supplementary Fig 4b). When investigating the average pup number born per litter, female FVB and B6.*Fancm^Δ^*^2/^*^Δ^*^2^ mice produced less litters, which were also smaller (Supplementary Fig 4c and d), suggesting that these mice could be useful models for the investigation of premature ovarian insufficiency seen in humans with biallelic mutations in *FANCM* [10]. This aside, the fertility of the mice provided an opportunity to use genetic approaches to study meiotic outcomes in a *Fancm^Δ^*^2/^*^Δ^*^2^ mammal.

### *Fancm* limits meiotic crossovers in mammals

We characterised the effect of *Fancm* loss on meiotic crossing over in F1 hybrids using three different methods. We began with male F1 mice, which were backcrossed to the FVB/N strain to generate BC1F1 offspring (Fig. 1a). Using a PCR-based genotyping method with a set of 271 informative SNP markers across 56 mice (n = 31 *Fancm^Δ^*^2/^*^Δ^*^2^, n = 25 *Fancm*^+/+^), we determined that F1.*Fancm^Δ^*^2/^*^Δ^*^2^ have a significant increase in crossover numbers and therefore total genetic map length. The genetic maps for *Fancm^Δ^*^2/^*^Δ^*^2^ and *Fancm*^+^*^/^*^+^ were 1,305 and 1,030 cM, respectively (p = 0.003, permutation test (B = 3,000)) (Fig. 1b.iii).

Using the same breeding strategy (Fig. 1a), we extended the crossover analysis with high-throughput “bulk” DNA sequencing with BC1F1 individual offspring and generated DNA sequencing datasets with 1-5X whole-genome coverage for each BC1F1 pup (total n = 372). Approximately four million informative SNP markers between C57BL/6J and FVB/N and were assayed to detect the strain origins in the BC1F1 genomes to call crossovers. With increased marker density, more crossovers were detected in the wild-type mice, and we observed an increase in total genetic distances in *Fancm^Δ^*^2/^*^Δ^*^2^ compared to *Fancm*^+/+^ and *Fancm*^+/^*^Δ^*^2^ pooled together (p = 0.03, permutation test (B = 1,000) with 1,339cM, 1,249 cM respectively)(Fig. 1b.ii), and the hyper-crossover phenotype due to the loss of *Fancm* was recessive (p = 0.45, permutation test (B = 1,000), 1,242 cM *Fancm*^+/^*^Δ^*^2^, 1,256cM *Fancm*^+/+^). Female genetic distances were higher in *Fancm^Δ^*^2/^*^Δ^*^2^ compared to *Fancm*^+/+^ and *Fancm*^+/^*^Δ^*^2^ but not significantly different (p = 0.15, permutation test (B = 1,000), 1,438 cM, 1,385 cM )(Fig. 1b.iv) and crossover distributions were unaffected (Supplementary Fig. 5).

Next, we assayed crossover frequency and distributions with single-gamete (haploid) sequencing in F1 males. We used a droplet-based method of partitioning individual gametes coupled with low level whole-genome amplification (Materials and methods). Total map lengths per sample within genotype groups were reproducible (Fig. 1b.i, Supplementary Fig. 6a, b) and no bias in marker segregation was observed (Supplementary Fig. 6c, d). Crossover frequencies across the genome were reproducible between single-gamete and bulk sequencing assays (Fig. 1c, Supplementary Fig. 7). Further, the total map lengths of all chromosomes for all conditions had a minimum approximate length of 50 cM, similar to the BC1F1 sequencing data (Supplementary Fig. 6b). These observations suggest that single-gamete sequencing can be reliably used for crossover detection. Differences in genetic distances along chromosomes in bins of 10 Mb were tested in genotype groups but no significant differences were observed for both assays for any one interval (Fig. 1c). To call crossovers using single-sperm sequencing data and downstream analyses, we developed an open-source bioinformatic tool kit containing *sgcocaller* and *comapr* [76] (see Materials and Methods). Consistent with the bulk sequencing and PCR-based methods, an increase in total crossover frequency and map length was observed in *Fancm^Δ^*^2/^*^Δ^*^2^ compared to *Fancm*^+/+^ (p = 0.001, permutation test (B = 1,000), 1,347 cM and 1,178 cM, respectively)(Fig. 1b.i). In conclusion, the triangulation of complementary and independent methods of crossover detection all support the finding that FANCM limits crossover formation in mammals, with the complementary strategies revealing an effect of a similar order of magnitude when considering sampling variance, and that marker densities vary substantially between the techniques used.

While genome-wide crossover numbers were elevated in *Fancm*-deficient mice, the general crossover distribution in *Fancm^Δ^*^2/^*^Δ^*^2^ appeared to be comparable to that of the wild type. The regions that had an increase of crossover frequency did not have a detectable unifying genomic feature (Supplementary Fig. 8-10). Nevertheless, the data from individual sperm show a significant increase in total crossovers per chromosome that was most notable for chromosomes 1, 4, 8 and 11 (Fig. 1) (one-sided t-test with multiple testing correction “fdr” with a total number of 19 tests). Crossover frequencies were also converted to genetic distances in centiMorgans per chromosome. The differences in total centiMorgans between genotypes per chromosome were tested using permutation-based methods, which showed a significant increase in *Fancm^Δ^*^2/^*^Δ^*^2^ for chromosomes 1, 2, 4, 8, 9, 11, 13, 18 before multiple testing correction (permutation p<0.05, B = 1,000), while chromosomes 1, 2, 4, 8, 11, 13 remained significant after multiple testing correction (using “fdr” for a total number of 19 tests) (Fig. 1d). An increase in crossover frequency on chromosome 11 in sperm from *Fancm^Δ^*^2/^*^Δ^*^2^ mice was detected with all three methods and an attempt to understand this phenomenon was pursued. Previous studies have highlighted how chromosome 11 has a particularly high crossover density per megabase in the wild type and have developed predictive models of crossover distributions [29, 77]. A linear regression model was fitted to our observed data using public data [77] as described [29] that used the physical chromosome size, meiotic DSB frequency and replication speed to predict crossover density. The predicted crossover densities were compared with observed crossover densities from this study. The model – that was developed for C57BL/6 x CAST – predicts lower crossover densities generally than are observed for the C57BL/6 x FVB/N data (this study), and predictions were better for *Fancm^Δ^*^2/^*^Δ^*^2^ than *Fancm*^+/+^. Chromosome 11 fit relatively well with the model in *Fancm^Δ^*^2/^*^Δ^*^2^ in both the pedigree- and sperm-based approaches with a much higher crossover density (Supplementary Fig. 11) revealing that chromosome size, DSB number and replication speed are good chromosome-scale crossover predictors for in *Fancm*-deficient mice. We interpreted this fit to indicate that the extra crossovers in *Fancm* loss of function mice had a relative distribution between chromosomes comparable to *Fancm*^+/+^.

During the development of our single-sperm sequencing and crossover mapping methods we attempted to identify crossovers with single-sperm RNA-seq data. In spite of careful data generation and analysis, however, this was unsuccessful. Cytoplasmic sharing of sperm is documented [78], but we hoped that a sufficient proportion of non-shared transcripts may be used for crossover mapping. We sorted haploid sperm cells from F1.*Fancm^Δ^*^2/^*^Δ^*^2^ and F1.*Fancm*^+/+^ to use as input for library preparation with the 10X Genomics single-cell gene expression protocol (Single Cell 3*^1^* v2). Cellranger (cellranger-mm10-3.0.0, GRCm38, annotation version 93) called over 15,000 barcodes per input sample as highconfidence single cells. However, even after rigorous quality control of called cells using best-practice workflows [79], we observed frequent and extensive regions of heterozygosity (variant allele frequencies 0.5) for putative haploid cells. The extent of heterozygosity was too high to allow mapping of crossovers from droplets containing one spermatid. The frequent detection of heterozygosity suggests that the degree of cytoplasmic transcript-sharing levels between spermatids prevents the accurate haplotyping of gametes that are needed for crossover mapping. Nevertheless, we could reproduce previous findings [80] that there exists a subset of transcripts that are more informative than average as to the nuclear genotype of a spermatid (Supplementary Fig. 12). Taken together these results suggest that “single-cell” sequencing of RNA from mouse sperm should be considered as a mixture of spermatogonial transcripts that are partitioned into the products of meiosis, and that do not necessarily represent the nuclear genome of post-meiotic sperm cells.

**Fig. 2:**
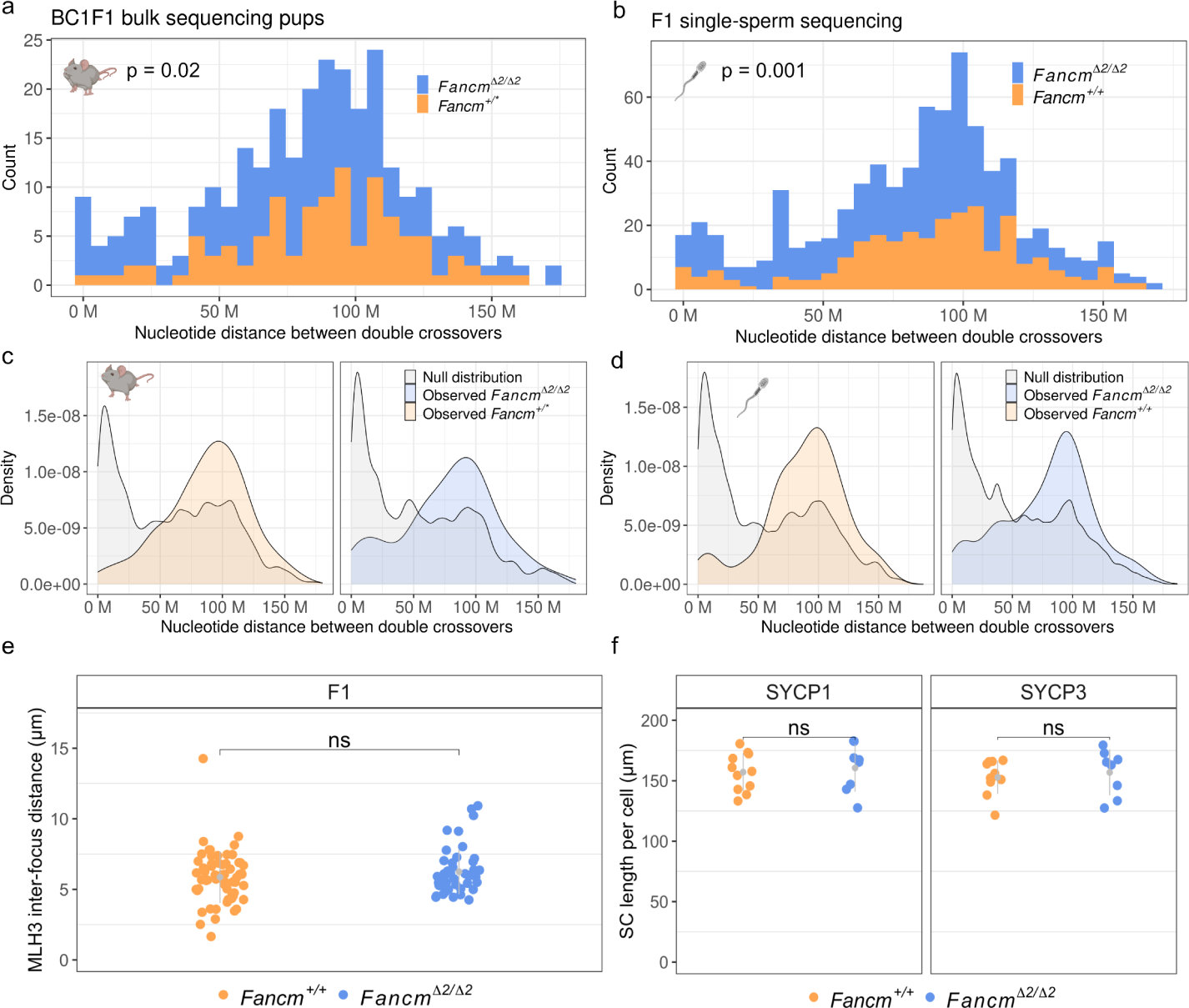
Extra crossovers in *Fancm* are consistent with the characteristics of class II crossovers. Interference is detected in *Fancm*-deficient male mice, but extra crossovers fit with a more random distribution and are not labelled by MLH1/3. a,b) Inter-crossover distances were quantified in the wild type and the mutant from the BC1F1-derived bulk sequencing and sperm sequencing data by only including chromosomes with exactly two crossovers. Printed p values were from Wilcoxon rank sum test. a, n = (175 *Fancm^Δ^*^2/^*^Δ^*^2^, 109 *Fancm*^+/*^, double crossovers per group. *Fancm*^+/*^ indicates pooling of *Fancm*^+/+^ and *Fancm*^+/^*^Δ^*^2^ data as their crossover rates were shown to not be significantly different.) b, n = (453 *Fancm^Δ^*^2/^*^Δ^*^2^,276 *Fancm*^+/+^). c,d) Crossover interference is observed in the wild type and the mutant as the observed inter-crossover distributions are more evenly spaced than an expected random distribution generated via permuting genotype group labels (B = 1,000). e) Cytological measurement of interference used MLH3 inter-focus distances in both the wild type and the mutant. n = 2 F1 (FVB/N x C57BL/6J) mice per genotype. Total cell analysed: 62 F1.*Fancm*^+/+^, 47 F1.*Fancm^Δ^*^2/^*^Δ^*^2^. f) Synaptonemal complex length was assessed by measuring SYCP1 and SYCP3 length in *µ*m per nucleus in pachytene spermatocytes. Total cells analysed: 16 F1.*Fancm*^+/+^ and 22 F1.*Fancm^Δ^*^2/^*^Δ^*^2^.

### Crossover interference and the obligate crossover are unaffected by a hyper-recombination phenotype

As we observed an increase in crossover frequency in *Fancm*-deficient mice, we sought to test how these extra crossovers are generated and if they show characteristics of the class I and/or class II pathways. Crossovers from the class I pathway are typically sensitive to crossover interference, meaning that they are more distantly spaced than would occur by chance. Therefore, we analysed spacing between two crossover events on the same chromosome. These intercrossover distances from our genetic data revealed that crossovers were more distantly spaced than random in all datasets analysed where sample size permitted (Fig. 2a,b) for both the wild type and the mutant. The random distributions of inter-crossover distances were generated through permutation via label swapping (Fig. 2c,d, Supplementary methods).

Given that there is an increase in crossover number in the *Fancm*-deficient mice, we tested if the proportion of class I and class II crossovers was affected. In the BC1F1 bulk sequencing experiment the distribution of inter-crossover distances was significantly different between the mutant and the wild type (p = 0.02, Wilcoxon rank sum test with continuity correction, n = (175,109) double crossovers per group) with the difference being attributable to a portion of closer double crossovers (Fig. 2a). The increased proportion of physically closer double crossovers in *Fancm*-deficient mice is consistent with the idea that there was an increase in class II crossovers. The same pattern was observed in the single-gamete sequencing data (p = 0.06, Wilcoxon rank sum test with continuity correction, n = (453, 276) double crossovers per group )(Fig. 2b). We used a complementary approach to test for characteristics of class I and class II crossovers in the *Fancm*-deficient mice by analysing MLH1 and MLH3 foci distributions, which are required for, and localise to, the sites of class I crossovers [51, 81–85]. Using labelling of MLH3 and MLH1 as a proxy for class I crossovers revealed no difference in event spacing and that crossover interference is present in the wild type and the mutant (Fig. 2e). The length of the synaptonemal complex is positively correlated with the number of class I crossovers [47–50]. Therefore we measured the length of the synaptonemal complex in the absence of *Fancm* by measuring the central and axial element, SYCP1 and SYCP3 respectively. Synaptonemal complex length appeared unchanged in pachytene in the mutant compared to the wild type (Fig. 2f). No differences in prophase I progression were observed in F1.*Fancm^Δ^*^2/^*^Δ^*^2^ compared to the wild type, with all meiotic prophase sub-stages from leptotene to diakinesis being observed in similar proportions (Supplementary table 1). These genetic and cytological data are consistent with an increase in class II non-interfering crossovers, but an unchanged class I crossover number and distribution.

**Table 1:** Offspring of B6.*Fancm^Δ^*^2/+^ x B6.*Fancm^Δ^*^2/+^ intercrosses. indicates p-value *<* 0.05, *** indicates p-value *<* 0.0005 (chi-square test).

|  | Male |  | Female |  |
| --- | --- | --- | --- | --- |
|  | Observed | Expected | Observed | Expected |
| *Fancm*<sup>+/+</sup> | 64 | 57 | 63 | 57 |
| *Fancm*<sup>+/<sup>Δ2</sup></sup> | 137 | 114 | 118 | 114 |
| *Fancm*<sup>Δ2/<sup>Δ2</sup></sup> | 45* | 57 | 29*** | 57 |
| Total | 246 |  | 210 |  |

As F1.*Fancm^Δ^*^2/^*^Δ^*^2^ males have increased crossover rates, we sought to cytologically investigate markers of DSB formation and repair in mouse spermatocytes using chromosome spreads. In order to analyse DSB formation we quantified RAD51 foci throughout early meiotic prophase I (Fig. 3a, b). The progressive decrease in variation and number of RAD51 focus counts in the wild type was consistent with previous work [21]. In *Fancm^Δ^*^2/^*^Δ^*^2^ spermatocytes, we found that RAD51 foci were slightly increased at pachytene in the mutant (p < 0.001) in both the F1 hybrid and the FVB/N inbreds, suggesting a delay in processing of joint molecules that are processed in the absence of *Fancm* (Fig. 3b). Elevated RAD51 at pachytene was also observed in *Fancm* homozygotes previously [6]. In order to assess if the elevated RAD51 foci numbers may contribute to DSB repair failure, we analysed *γ*H2AX localisation. Using *γ*H2AX, sex body formation appeared normal in F1.*Fancm^Δ^*^2/^*^Δ^*^2^, and no persistent *γ*H2AX was observed on autosomes at pachytene using meiotic chromosome spreads. We next sought to analyse DSB repair and pathway choice. We analysed the loading of the ZMM MSH4, which initially localised to a majority of recombination sites, and along with its binding partner MSH5, is required for normal synapsis and crossover formation [86–88]. MSH4 foci numbers were unchanged between *Fancm* genotypes in recombination intermediates labeled by MSH4 (Fig. 3a). Similarly, MLH1 and MLH3 foci numbers, markers of class I crossovers, were unchanged in *Fancm*deficient mice (Fig. 3a,c). We analysed if the extra crossovers in *Fancm^Δ^*^2/^*^Δ^*^2^ affect meiotic completion. The obligate crossover was unaffected in *Fancm^Δ^*^2/^*^Δ^*^2^ as bivalent formation was unchanged when analysed with Giemsa-stained male meiotic metaphase I chromosomes (Fig. 3d,e). Similar quantification of meiotic metaphase I on histological sections revealed no detectable difference in unbalanced univalents between genotypes (Supplementary table 1). Taken together, these data from different steps of DSB repair and crossover formation suggest that the extra crossovers in a mammalian *Fancm* mutant are not class I crossovers and do not dramatically affect meiotic progression.

**Fig. 3:**
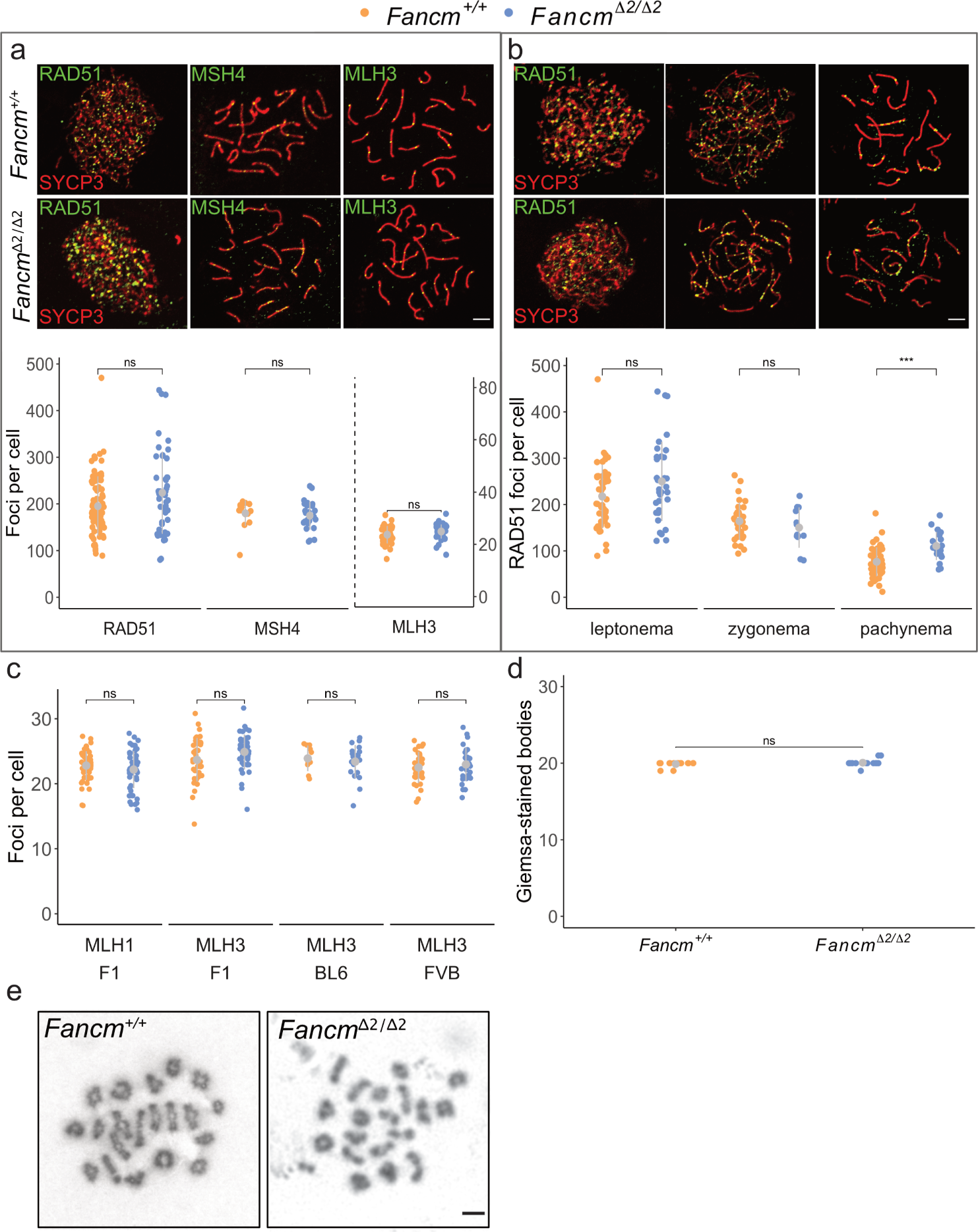
Meiotic double-strand break repair was relatively normal in *Fancm* *^Δ^*^2/^*^Δ^*^2^ spermatocytes. a) Representative images of F1.*Fancm*^+/+^ and F1.*Fancm^Δ^*^2/^*^Δ^*^2^ spermatocyte chromosome spreads from meiotic prophase, stained for the indicated proteins (above) and quantification of foci per nucleus in wild-type and *Fancm^Δ^*^2/^*^Δ^*^2^ spermatocytes. Each data point is a count from one cell. Scale bar is 5 *µ*m. Bars indicate mean *±* s.d. F1.*Fancm*^+/+^: RAD51 foci at leptonema (217.65 *±* 68.8), MSH4 foci at zygonema (164.2 *±* 40.4) and MLH3 foci at pachynema (23.7 *±* 3.3). F1.*Fancm^Δ^*^2/^*^Δ^*^2^: RAD51 foci at leptonema (250.0 *±* 85.5), MSH4 foci at zygonema (150.3 *±* 43.6) and MLH3 foci at pachynema (24.9 *±* 2.7). Total RAD51-labelled cells analysed: 79 F1.*Fancm*^+/+^, 46 F1.*Fancm^Δ^*^2/^*^Δ^*^2^. Total MSH4-labelled cells analysed: 16 F1.*Fancm*^+/+^ and 26 F1.*Fancm^Δ^*^2/^*^Δ^*^2^. Total MLH3-labelled cells analysed: 40 F1.*Fancm*^+/+^ and 47 F1.*Fancm^Δ^*^2/^*^Δ^*^2^. b) RAD51 foci numbers are higher at pachynema in *Fancm^Δ^*^2/^*^Δ^*^2^. Grey bars indicate mean *±* s.d. Each data point indicates a count from one nucleus. F1.*Fancm*^+/+^ RAD51 foci at leptonema (217.7 *±* 68.8), zygonema (164.2 *±* 40.4) and pachynema (76.5). *Fancm^Δ^*^2/^*^Δ^*^2^ RAD51 foci at leptonema (250.0 *±* 85.5), zygonema (150.3 *±* 43.6) and pachynema (110.6 *±* 30.3). Scale bar is 5 *µ*m. (*** indicates p-value *≤* 0.001 (unpaired t-test). 2 F1 mice analysed per genotype at each meiotic phase.) c) Variation of MLH3 foci in B6, FVB, and F1 backgrounds. 89 F1.*Fancm*^+/+^, 101 F1.*Fancm^Δ^*^2/^*^Δ^*^2^, 10 B6.*Fancm*^+/+^, 20 B6.*Fancm^Δ^*^2/^*^Δ^*^2^, 32 FVB.*Fancm*^+/+^ and 32 FVB.*Fancm^Δ^*^2/^*^Δ^*^2^ cells were analysed. Scale bar is 5 *µ*m. d) Representative meiotic metaphase I spreads for B6.*Fancm*^+/+^ and B6.*Fancm^Δ^*^2/^*^Δ^*^2^. e) The number of Giemsa-stained bodies per spermatocyte in *Fancm*^+/+^ and *Fancm^Δ^*^2/^*^Δ^*^2^. 31 *Fancm*^+/+^ and 19 *Fancm^Δ^*^2/^*^Δ^*^2^ cells were analysed.

### Spermatogenesis and oogenesis are perturbed in *Fancm^Δ^*^2/^*^Δ^*^2^ mice

One of the few unifying phenotypes of mouse models from the Fanconi anaemia pathway, including *Fancm*-deficient mice [5,6,8], is abnormal gonad development and sub-fertility [89]. We explored this further and found that *Fancm^Δ^*^2/^*^Δ^*^2^ testicular weight was reduced (Fig. 4a,b). The reduction in testes weight was significant in all strains: FVB.*Fancm*, B6.*Fancm* and F1.*Fancm* (Fig. 4b). The reduction in the F1.*Fancm* however, was not as strong as the reduction in testicular weight in the two inbred strains. In all three strains, daily sperm production was reduced in the mutant indicating that *Fancm* is required for normal sperm production levels (Fig. 4c). Further, computer-assisted sperm analysis revealed that for those sperm that were produced, both B6.*Fancm* and FVB.*Fancm* had a decrease in the percentage of motile and progressively motile sperm (Fig. 4d,e), however this only resulted in reduced litter sizes in B6.*Fancm*. In females, ovary weights were reduced in B6.*Fancm*; however, the difference was less dramatic than what was seen for male gonads. FVB.*Fancm* and F1.*Fancm* ovary weights were lower than their wild-type litter mates on average but did not reach statistical significance (Fig. 4f). However, a dramatic reduction in the number of follicles per ovary was observed in *Fancm*-deficient FVB/N, C57BL/6J and F1 mice, consistent with a premature ovarian insufficiency phenotype (Supplementary Fig. 13, Supplementary Fig. 4). Further, super-ovulated F1 mutants had a reduction in oocyte numbers (Fig. 4g).

**Fig. 4:**
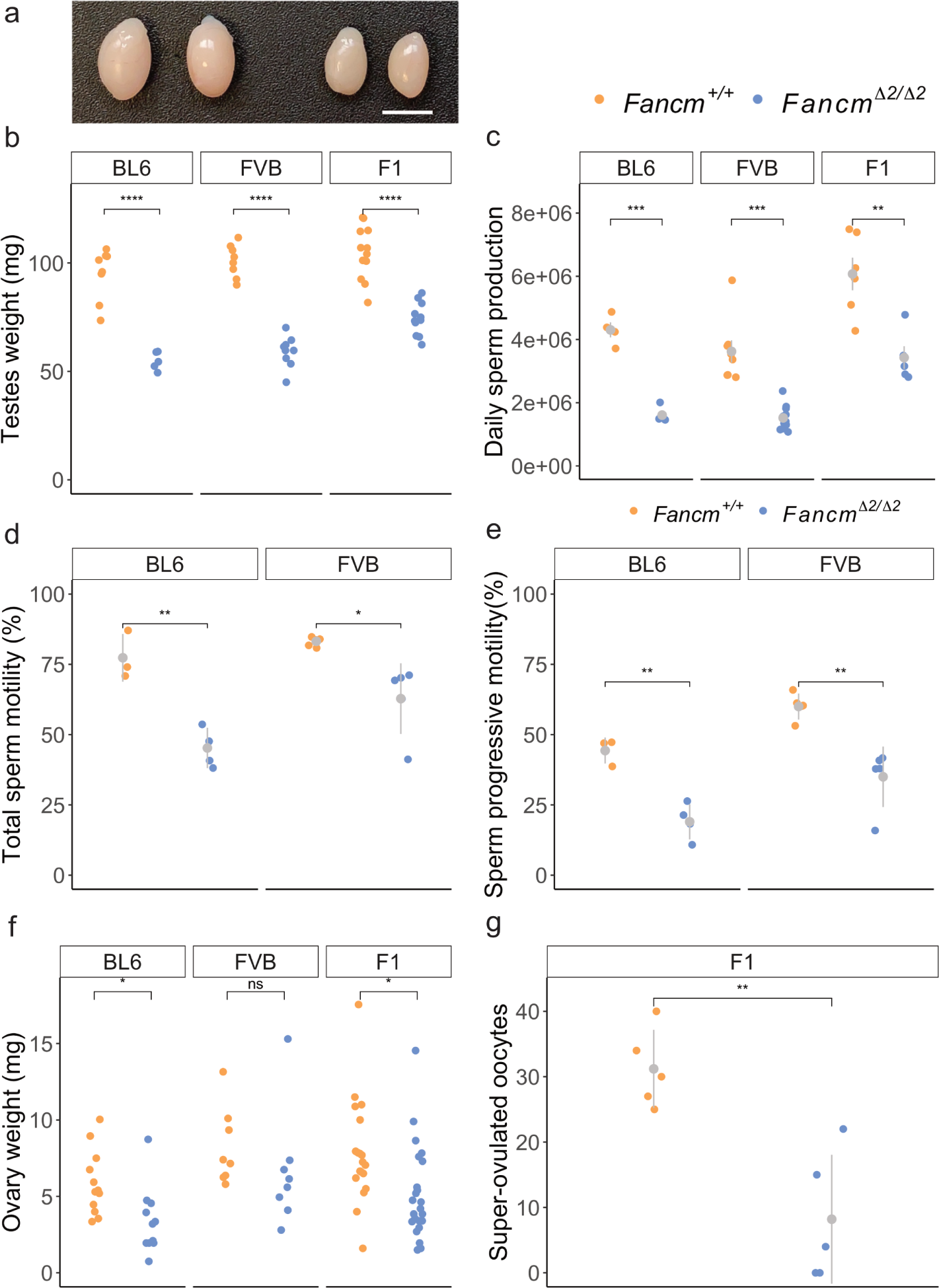
Hypogonadism and gametogenesis defects in both sexes of *Fancm* -deficient mice. a) Testes from B6.*Fancm*^+/+^ (left) and B6.*Fancm^Δ^*^2/^*^Δ^*^2^ (right) mice. Scale bar = 5 mm. b) 6 to 14 week old testes weights were lower in the mutants in all strains. c) Quantification of daily sperm production. d) Total sperm motility, and e) sperm progressive motility was assessed with computer-assisted sperm analysis from 10 week old mice. f) Ovary weights were quantified across the three strains used from 5 week orPoaldgeer1m4icoef. ^7^g0) F1 females were super-ovulated and *Fancm*-deficient mice produced less oocytes. 5 mice per genotype. * indicates p *≤* 0.05, ** indicates p *≤* 0.01, *** indicates p *≤* 0.001 and *** indicates p *≤* 0.0001.

### cGAS-STING drives spermatogonial defects in *Fancm* mice

A Fancm point mutant mouse has previously been shown to be characterised by smaller testes due to impaired primordial germ cell proliferation, in addition to a progressive loss of male germ cells in adulthood [6]. To investigate the origin of the subfertility of our B6.*Fancm^Δ^*^2/^*^Δ^*^2^ males and the reduced daily sperm production across all Fancm mutant strains, we analysed the seminiferous tubules of wild-type and mutant mice. Histological analysis of the FVB.*Fancm^Δ^*^2/^*^Δ^*^2^ strain showed no overt differences in the testicular histology between mutant and wild type littermates (Supplementary Fig. 18). FVB.*Fancm^Δ^*^2/^*^Δ^*^2^ seminiferous tubules appeared replete with all germ cell types present, suggesting the strong reductions in testes weight and daily sperm production (Fig. 4b-e), are likely driven by deficient primordial germ cell proliferation as described previously [6]. Histological analysis of B6.*Fancm^Δ^*^2/^*^Δ^*^2^ however, showed a number of defects. Most notably, and consistent with the progressive loss of germ cells seen previously due to Fancm deficiency [6], a small but significant subset of seminiferous tubules were characterized by either partial or complete absence of germ cells from the epithelium (Sertoli cell only, Fig. 5a,b, Supplementary Figs. 14, 18). In the least severely affected of these, vacuoles between adjacent Sertoli cells were observed in addition to areas of Sertoli cell cytoplasm devoid of germ cells (Fig. 5b). In more moderately affected tubules whole populations of specific germ cell types were depleted (Fig. 5b, Supplementary Fig. 14). The most severely affected tubules were characterized by a Sertoli cell-only phenotype, wherein the loss of integrity of the seminiferous epithelium ultimately led to the depolarisation of Sertoli cells from the basement membrane and their collapse into multi-nucleated symplasts (Fig. 5a,b, Supplementary Fig. 14).

**Fig. 5:**
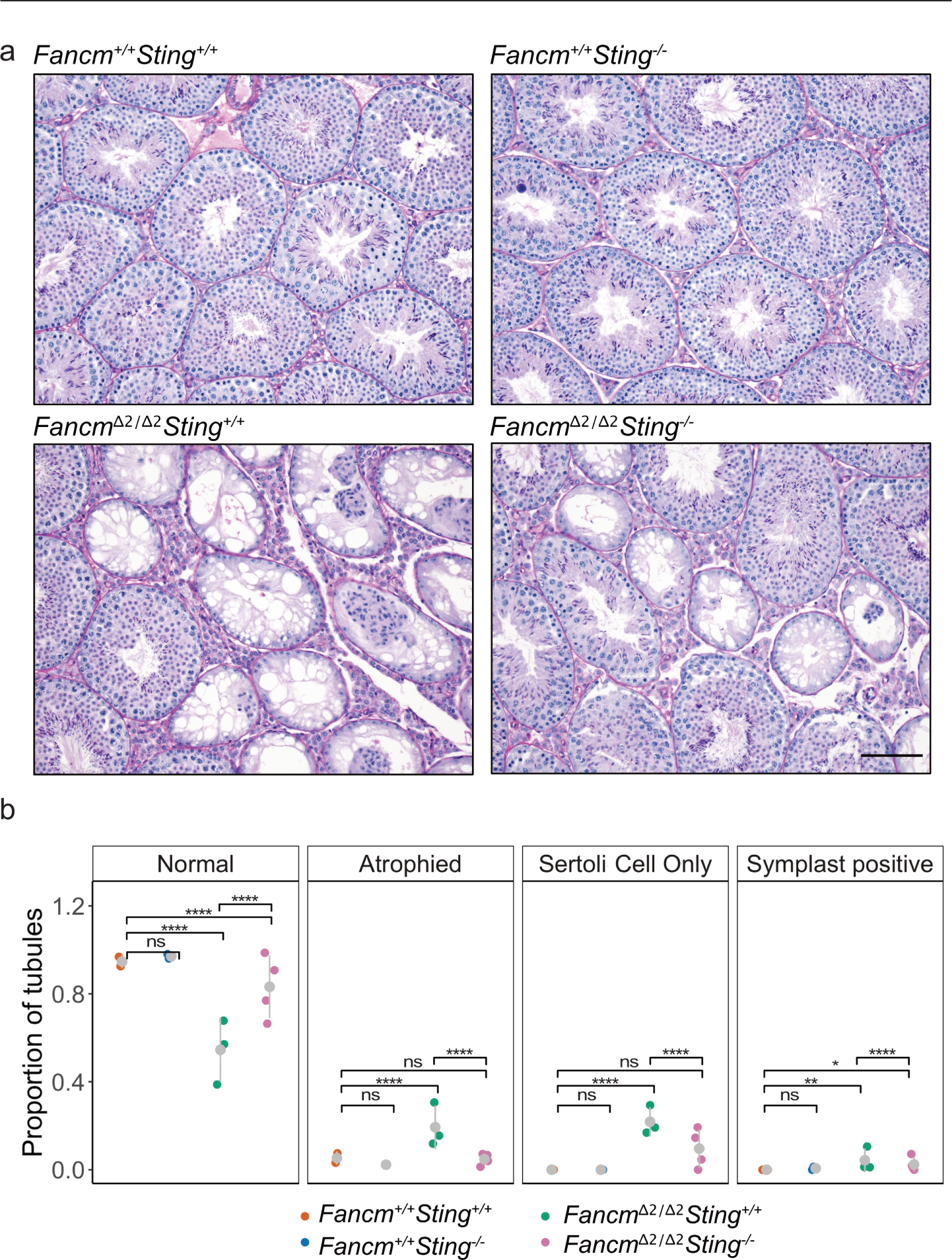
*Sting* depletion partially rescues histological defects in the seminiferous tubules in *Fancm* -deficient mice. a) H and E stain of seminiferous tubules in B6.*Fancm*^+/+^ and B6.*Fancm^Δ^*^2/^*^Δ^*^2^ mice. Scale bar = 100 *µ*m. Wild-type tubules are relatively replete in their appearance. Mutant mice have a heterogeneous reduction in spermatocytes (hypospermatogenesis) at the level of individual tubules. A mixture of normal, atrophied, symplast-positive and Sertoli cell-only tubules were be observed. *Fancm^Δ^*^2/^*^Δ^*^2^*Sting* ^-/-^ testes have more full tubules than the *Fancm* single mutant (p-adjust < 0.0001, b) Quantification of the PAS-stained seminiferous tubule cross-sections. Each data point represents data from one mouse shown as a proportion for the given classi_P_fic_a_a_g_t_e_io_1_n_7_. _o_*_f_*^7^*^0^* indicates p-value *≤* 0.0001, ** indicates p-value *≤* 0.01, * indicates p-value *≤* 0.05 (pair-wised proportion test with multiple testing correction “fdr”). A total of 290 *Fancm*^+/+^*Sting* ^+/+^, 305 *Fancm*^+/+^*Sting* ^-/-^, 479 *Fancm^Δ^*^2/^*^Δ^*^2^*Sting* ^+/+^, 605 *Fancm^Δ^*^2/^*^Δ^*^2^*Sting* ^-/-^ tubules were analysed.

An initial analysis of apoptosis with histological sections from C57BL/6J and FVB/N testes sections was performed by immunolabelling cleaved caspase-3 and -7 (Supplementary Fig. 15). We did not detect a change in the number of caspase3/7 positive germ cells per seminferous tubule in the mutants compared to the wild type. More refined analysis of the B6.*Fancm^Δ^*^2/^*^Δ^*^2^ seminiferous epithelium suggested that there were at least two possible points of reduced daily sperm production, in addition to the well documented reduced proliferation of primordial germ cells via the p21-p53-ATM axis [6]. As shown in Supplementary Figure 14b, in the B6.*Fancm^Δ^*^2/^*^Δ^*^2^ seminiferous tubules characterised by progressive germ cell loss, pachytene spermatocytes often appeared to be the first germ cell population to be absent. Stage V-VIII tubules that harboured their characteristic populations of spermatids, spermatogonia and other spermatocyte types [90], were often either partially or completely devoid of pachytene spermatocytes (Xsquared = 19.314, df = 1, p < 0.001, Supplementary Fig. 16, Supplementary Table 5). This phenotype appeared to be driven by a significant increase in the death of spermatocytes at mid-pachytene. Indeed, analysis of stage V seminiferous tubules revealed a significant increase in the rate of pachytene spermatocytes with PAS-positive cytoplasm – PAS is suggested to stain apoptotic cells [91] – in B6.*Fancm^Δ^*^2/^*^Δ^*^2^ testes compared to wild type controls (p < 0.05, Supplementary Fig. 16f).

A second point of loss may be at the first meiotic division in metaphase I-anaphase I. At stage XII of spermatogenesis when the meiotic divisions are normally detectable on histological sections, the number of cells in metaphase I-anaphase I was comparable between B6.*Fancm^Δ^*^2/^*^Δ^*^2^ and littermate controls (Supplementary Fig 16c). The number of metaphase I-anaphase I spermatocytes that remained into the succeeding stage I, wherein meiosis is normally complete, was also comparable between genotypes (Supplementary Fig 16c). Alignment of bivalents on the metaphase plate and their segregation also appeared unchanged (Fig. 3, Supplementary Fig 16c). However, a strong and significant increase in PAS-positive metaphase I-anaphase I spermatocytes, indicative of meiosis arrest, was observed in stage XII tubules (Supplementary Fig. 16). Likewise, a more than two-fold increase in PAS-positive metaphase I-anaphase I or pyknotic cells was observed in stage I tubules (Welch Two Sample t-test, p < 0.0001, n = four mice per genotype, Supplementary Fig. 16e). However, no abnormalities were detected with neither SYCP3 staining meiotic prophase sub-stages, nor two dimensional metaphase I chromosome spreads (Fig. 3, see Discussion). Histological analysis of spermatogenesis detected occasional binucleated round spermatids indicative of failed cytokinesis in B6.*Fancm^Δ^*^2/^*^Δ^*^2^ but not in wild type controls. In conclusion, within B6.*Fancm^Δ^*^2/^*^Δ^*^2^ mice, the loss of FANCM resulted in at least three detectable defects using histology. This aside, sperm were produced and mice retained some level of fertility.

We next analysed the innate immune response as previous studies have revealed that genomic instability in mouse models can also activate cGAS-STING pathway [92, 93], which is a part of the innate immune system that detects cytosolic DNA [94–96]. We sought to investigate whether the cGAS-STING pathway could be driving the reduction in germ cells in the B6.*Fancm^Δ^*^2/^*^Δ^*^2^ mice. *Fancm Sting* double mutants, compared to *Fancm* single mutants, had an increase of approximately 20% in testicular weight, while epididymis weight and daily sperm production were unchanged, and far inferior to wild-type levels (Supplementary Fig. 17). Next we performed histological analyses with the seminiferous tubules. This revealed that the seminiferous tubules of the *Fancm Sting* double mutants compared to *Fancm* had a statistically significant increase in seminiferous tubules containing complete spermatogenesis and histological observation of spermatogenesis compared to the *Fancm* single mutant (p < 0.0001, pair-wised proportion test after multiple testing correction) (Fig. 5, Supplementary Fig. 15). In contrast, folliculogenesis in *Fancm*-deficient mice was not ameliorated by the removal of the *Sting* pathway (Supplementary Fig. 17). These data support the hypothesis that the cGAS-STING pathway drives a portion of the attrition of male germ cells in the absence of the Fanconi anaemia pathway, highlighting that inhibitors of the cGAS-STING pathway may have therapeutic benefits for fertility in males with germline genomic instability conditions.

Given the interaction between *Fancm* and the innate immune response via the cGAS-STING pathway, we sought to test alternative inflammatory pathways. Previous work has revealed that there is a female-biased embryonic lethality in the absence of *Fancm* [7, 97]. During the course of our experiments we observed an under representation of B6.*Fancm^Δ^*^2/^*^Δ^*^2^ mice, which was more severe for females (Table 1). Previous studies have observed sub-Mendelian birth rates of female *Fancm*-deficient mice [5–7], however with a larger sample size in our study we identified this to occur for B6.*Fancm^Δ^*^2/^*^Δ^*^2^ males too. FVB.*Fancm^Δ^*^2/^*^Δ^*^2^ were not underrepresented in either sex however, and neither were F1 mutants descendent from a cross between B6.*Fancm*^+/^*^Δ^*^2^ and FVB.*Fancm*^+/^*^Δ^*^2^ ((Supplementary tables 1 and 1)). These data suggest that the sub-Mendelian birth rates of *Fancm* mutants in our experiments is specific to the C57BL/6J background and is a recessive trait. Previous observations of bias in lethality of female embryos was shown to be due to inflammation caused by genomic instability and could be alleviated by supplementing the drinking water of pregnant mothers with the anti-inflammatory drug ibuprofen [7]. We tested whether ibuprofen reversed the seminiferous tubule defects of *Fancm*-deficient mice, however there was no effect compared to controls (Supplementary Fig. 18).

## Discussion

### *Fancm* performs distinct functions during reproduction

*Fancm* is essential for efficient homologous recombination by directing the outcomes of repair intermediates. This study identifies *Fancm* being a rate-limiting step during mammalian meiotic crossover formation. This study also extends the understanding of the role of *Fancm* in fertility. We propose that these two functions are distinct, as meiotic outcomes appear relatively unaffected at a chromosomal and molecular scale, whereas gamete numbers are affected in males and females.

We suggest that the meiotic hyper-crossover and other spermatogenesis phenotypes that occur due to the loss of *Fancm* are separable for several reasons:

1. No notable defects in bivalent formation compared to the wild type that could account for such a dramatic reduction in spermatogenesis and the requisite stem cells. The meiotic program progresses normally when considering direct immunofluorescent readouts of the synaptonemal complex formation and meiotic DSB repair. There are more meiotic DSBs that persist at pachytene than in the wild type ( [6] and this study), but given the rate of daily sperm production in mutants and healthy pups that were born, it does not seem that these lead to major meiotic errors and perhaps simply reflect altered and later processing of a subset of meiotic DSBs; 2) the heterogeneous Sertoli cell-only phenotype of some seminiferous tubules may be due to stochastic DNA repair-related aberrations that occurred during the establishment of the germ cell lineages early in embryo development and perhaps also during failed maintenance of spermatogonia that only achieve a limited number of rounds of spermatogenesis, which would be consistent with other mice lacking the FA pathway [6, 89]. Similarly, some tubules have a “lacy” phenotype that may be due to the loss of individual spermatogonial stem cells. Phenotypes that are driven by strong meiotic DSB repair defects have a homogeneous arrest at the pachytene and typically produce no round spermatids or spermatozoa, when assessed with histological sections of the seminiferous tubules [98, 99]. No such pachytene arrest is observed in F1.*Fancm^Δ^*^2/^*^Δ^*^2^, neither with surface spreads that visualise the synaptonemal complex, nor with histological analysis. However, we do not exclude the possibility that the hyper-crossover *Fancm* could make a minor contribution to reproductive defects; and
2. previous work has shown a reduction in primordial germ cells in the absence of *Fancm* [6], and an increase in specific mutational signatures [100], which may account for the reduction in gamete numbers and litter sizes. One possibility of how these two functions co-exist is through the vertebrate-specific addition of the ERCC4 domain. The translocase domain would be the domain that remodels DSB repair intermediates to limit crossover rates. However, the ERCC4 domain may be involved in FA pathway activation and in turn protects stem cells from damage. In summary, we propose that these observations support the hypothesis that the meiotic anti-crossover function of FANCM can be uncoupled from its role in promoting genomic stability and efficient spermatogenesis.

### FANCM is an evolutionarily conserved crossover-limiting factor

Here we demonstrate that *Fancm* limits meiotic crossovers in mammals. This is the first report of a species with a *bona fide* FA pathway has been shown to use FANCM to limit meiotic crossover rates. The limited shuffling of genetics enforced by *Fancm* is conserved across diverse eukaryotes including mammals (this study), plants, fission yeast and fly [69, 70, 73, 101, 102]. This raises the question of whether or not FANCM achieves its CO-limiting function with other components of the FA pathway, which will be interesting for future studies to discern. CO formation is a tightly regulated and well conserved process. DNA doublestrand breaks outnumber COs consistently across diverse species, which suggests that there are conserved DNA repair factors that keep CO numbers below their possible upper bounds. This study and previous studies highlight *Fancm* as a major driver of this phenomenon [69,70,73,101,102]. Why crossover numbers per cell are limited is difficult to discern. One hypothesis is that a baseline level of crossovers at meiosis and therefore only once per generation has a net positive effect for a population in a changing environment. However, there will also be likely negative consequences for a number of individuals associated with the decoupling of favourable combinations of linked alleles that have been selected for on evolutionary timescales [103]; therefore, on average parental combinations of alleles could be more beneficial than excessive shuffling of allelic combinations. Another hypothesis is that excessive crossovers impede correct segregation of chromosomes however this appears unlikely as a universal rule given that significant increases in crossover frequency did not affect chromosome segregation in: this study; mutants of Arabidopsis; or Drosophila mutants [69, 74, 101]. Also significant differences in crossover rates are normal within species, such as differences in crossover rates between males and females [104, 105], and some species naturally have very high levels of crossovers per chromosome [45].

Several crossover-limiting factors have been identified, many of which play important roles in maintaining genomic stability [46, 67, 68, 74, 106]. The molecular function of FANCM is consistent with an anti-crossover factor as it can act on a diverse range of branched DNA substrates, many of which are produced during meiotic DSB repair such as displacement loops and Holliday junctions that could progress to crossover formation. Therefore, the loss of FANCM function would force the repair of joint molecules to occur using alternate repair pathways and non-crossover outcomes. All studies of the anti-crossover role of FANCM have characterised loss of function alleles and *Fancm* has not been identified in studies that screened for natural variants that influence crossover rates to the best of our knowledge. Other meiotic crossover-limiting factors that tend to only be characterised as knockouts are Sgs1/BLM, RTEL-1 homologues and their binding partners [61, 68, 69, 74, 107–109], which are also required for genomic stability [110–113]. Perhaps the loss of function phenotypes are only identified in laboratory settings as their somatic functions tolerate little to no variation, which can be seen in their respective chronic paediatric diseases Fanconi anaemia, Bloom syndrome and Dykeratosis congenita. This contrasts with natural alleles of genes such as PRDM9, RNF212, HEI10, REC8 that are associated with the location of recombination hotspots, meiotic chromosome axis structure, and global crossover rates [24, 25, 114, 115].

The extra crossovers in F1.*Fancm^Δ^*^2/^*^Δ^*^2^ were not linked to overt changes with respect to relative distributions of crossovers along chromosomes. Further, these extra crossovers in *Fancm*-deficient mice are not labelled by MLH1/3 and are distributed with less even spacing than occurred in the wild type. This is consistent with an increase in non-interfering (class II) crossovers. These findings are consonant with what was observed in Arabidopsis [67]. A difference between Arabidopsis and mouse *Fancm* knockouts however, is that in the mouse, FANCM limits crossovers in inter-specifc F1 crosses; the strong anti-crossover activity of *AtFancm* is limited to inbreds or long runs of homozygosity [67,72,106]. We speculate that these differences in crossover formation in inter-specific hybrids may reflect alternate regulation of the early stages of strand invasion. The crossover density prediction model that we implemented from [29] suggests that physical chromosome size, replication speed and DSBs are good predictors of the extra crossovers that occur in the absence of *Fancm* (class II crossovers), just as they were good predictors of class I crossovers in the wild-type cross between C57BL/6 and CAST mice in [29]. Therefore, we propose that: 1) in mammals *Fancm* limits class II crossovers; and 2) while class I and class II crossovers have differing genetic requirements, class I and II crossover formation pathways have similar characteristics with respect to the genomic features that influence crossover formation.

### Reproductive defects occur at distinct periods of development in***Fancm* mice**

Humans and mice that lack proper *Fancm* function are infertile or have severely impacted reproductive capacity [5, 6, 8–10]. This is different from other species where loss of *Fancm* has no effect on gamete numbers despite leading to massive increases in total crossover numbers. These differences and other data that we present about meiotic outcomes further support the idea that the reproductive defects in the absence of *Fancm* are not tied to the anti-crossover role at meiosis.

The gonads of male and female *Fancm* knockouts were reduced in weight in two inbred strains and the F1 and is consistent with hypogonadism phenotypes in human Fanconi anaemia patients and other *Fanc* gene mouse models [5, 6, 8, 14, 89, 116]. Similarly, a consistent reduction in daily sperm production in *Fancm*-deficient mice was observed in all strains. Histological analyses of the testis revealed a dramatic phenotype in the C57BL/6J strain on seminiferous tubule composition; notably, a progression towards Sertoli cell-only phenotype. Direct analysis of markers of DSB formation and repair did not suggest a strong, or any, defect in meiotic DSB repair in the absence of *Fancm*. Similarly chromosome dynamics were unchanged when considering synapsis and chromosome segregation. Nevertheless, histological analyses showed unambiguous phenotypes when considering whole tubules, as opposed to individual meiocytes, such as complete absence of large groups of pachytene cells that would be found in comparable tubules in the wild type. Considering these data together, it seems most plausible that any recombination defect that may exist in *Fancm*-deficient mice is mild, particularly given that daily sperm production is measured in the millions and litter sizes from male mutants are relatively unchanged. However, an alternative interpretation could be that a particularly mild recombination defect triggers a checkpoint arrest, which is reflected by the PAS-positive metaphase I-anaphase I cells. In B6.*Fancm* we found that a portion of the defects in spermatogensis is driven by the cGAS/STING-dependent pathway, which raises the question as to whether a cGAS/STING inhibitor could offer the therapeutic benefit of improving fertility rates in males with genetic conditions rooted in genomic instability.

In contrast to B6.*Fancm*, for FVB.*Fancm* or F1.*Fancm*, a lesser effect was on gonad weight, daily sperm and seminiferous tubule composition was detected in experiments with histopathology. These data suggest that the B6.*Fancm* seminiferous tubule phenotype is recessive in the F1.*Fancm* and the causal variant(s) can likely be positionally cloned in future studies to identify modifiers of genomic instability phenotypes. The differing phenotypes of C57BL/6J and FVB/N seminiferous tubule phenotypes reveal a difference between the two strains that appears unlinked to *Fancm*, however the relevance of this is unclear and could be investigated in future studies.

In summary, despite the defects in gametogenesis (this study, and [5,6,8–10]) we show that *Fancm* knockouts are capable of producing offspring, which we could leverage to reveal the anti-crossover function of mammalian FANCM. The capacity of mouse knockouts to produce a tractable number of offspring has been shown for multiple *Fanc* gene knockout mice, highlighting the utility of these DNA damage response defective models to inform clinical fertility research for patients with chronic genetic conditions. Further, as human infertility can present before other serious symptoms caused by genetic conditions, there are opportunities to improve diagnostic pathways for life-limiting diseases by considering reproductive status [89, 117].

## Materials and methods

### Ethics statement

All animal procedures were approved by the Animal Ethics Committees at St Vincent’s Hospital Melbourne and Monash University and conducted in accordance with Australian NHMRC Guidelines on Ethics in Animal Experimentation.

### Mouse generation and genotyping

The *Fancm^Δ^*^2/^*^Δ^*^2^ mice were generated independently in FVB/N and C57BL/6 backgrounds. The same guide RNAs and Cas9 were used to target exon 2 of *Fancm*. For genotyping, wild-type and mutant alleles were amplified with oligos forward primer (5*^1^* CGGGGCGGAATGCTAAACTT 3*^1^*), reverse primer 1 (5*^1^* ACACACAGGGACAGAGAACACTC 3*^1^*, spans deleted region) and reverse primer 2 (5*^1^* GGAAAAGAGAAAAGAAAAGGGGGA 3*^1^*) and produced bands of 311 bp and 279 bp respectively that were visualised on a 2% agarose gel. *Sting^-/-^* mice have been described previously [118] and the *Sting* ^-/-^ mice were backcrossed for at least 10 generations on a C57BL/6 background by [119]. The oligos used for genotyping were forward primer 1 (5*^1^* GCTGGGAATTGAACGTAGGA 3*^1^*), forward primer 2 (5*^1^* GTGCCCAGTCATAGCCGAAT 3*^1^*, spans deleted region) and reverse primer (5*^1^* GAGGAGACAAAGGCAAGCAC 3*^1^*).

### Genomic DNA preparation and sequencing

Genomic DNA was extracted using the phenol-chloroform method. Briefly, a 0.5 mm of tail was placed in a 1.5 mL microfuge tube with 500 *µ*L of tissue digestion buffer (100 mM Tris-Cl pH 8.0, 5 mM EDTA pH 8.0, 200 mM NaCl, proteinase K 0.4 mg/mL) overnight on a shaking platform at 55*^◦^*C. The next day, 1 *µ*L of 10 mg/mL RNAse A was added to each sample and incubated at 55*^◦^*C for 30 min. 0.7 mL of neutralized phenol/chloroform/iso-amyl alcohol (25:24:1) was added to each sample and mixed vigorously at 4*^◦^*C on a clinical rotor for 1 hour. Samples were spun in a bench top centrifuge at max speed (21,000 x g) for 5 min and the upper aqueous phase containing DNA was transferred to a new 1.5 mL microfuge tube containing 1 mL of 100% ethanol to precipitate the DNA. The samples were spun at max speed for 5 min and supernatant was removed. DNA pellet was washed with cold 70% ethanol and spun again at max speed for 5 min. 1 X TE buffer was added to the samples to resuspend the DNA and placed on a 55*^◦^*C block to evaporate residual ethanol. All samples were checked for protein contamination by using Nanodrop A_260_/A_280_ and DNA quality by running samples on a 0.8% agarose gel to ensure there was no DNA shearing.

For bulk sequencing, genomic DNA was sent to BGI (China) or AGRF (Australia) and sequenced with DNBSeq or Illumina platforms respectively with between 1X and 5X coverage.

### Haploid nucleic isolation for droplet-based single cell library preparation

To obtain single haploid cells, testes were harvested from male mice. On ice, the tunica albuginea was removed to release the seminiferous tubules into a 1.5 mL microfuge tube and incubated on ice for 30 min. The spleen was also harvested and homogenised through a 40 *µ*m strainer to act as a diploid control for flow cytometry.

600 *µ*L of testes or 300 *µ*L of splenic homogenate was transferred to a new 1.5 mL LoBind microfuge tube. 1 mL of chilled Nuclei EZ Lysis buffer (Sigma) was added to each sample, inverted twice, and incubated on ice for 2 min. Samples were centrifuged at 500 x g for 5 min at 4*^◦^*C and supernatant was removed, ensuring that the pellet was still immersed. Samples were resuspended with 1 mL of Nuclei EZ Lysis buffer, inverted several times and incubated at 500 x g for 5 min at 4*^◦^*C. Samples were centrifuged at 500 x g for 5 min at 4*^◦^*C and supernatant was removed. 1 mL of Nucleic Wash and Resuspension Buffer (NWRB; PBS with 1% BSA) was added slowly to the samples to avoid disruption of the pellet and allow buffer interchange. Samples were centrifuged at 500 x g for 5 min at 4*^◦^*C and resuspended with 1 mL of NWRB, and this was repeated. Samples were centrifuged at 500 x g for 5 min at 4*^◦^*C and supernatant removed. Sample was resuspended with 1 mL NWRB with DAPI (10 *µ*g/mL). The samples were then filtered through using a 40 *µ*m Flowmi cell strainer into a 5 mL round bottom polystyrene tube.

Using the spleen sample as a control for the diploid peak, 100 000 haploid nuclei was sorted into a round-bottom 96-well plate with 50 *µ*L of NRWB + 0.4% BSA. Sorting was performed using an BD FACSARIA II cell sorter. After sorting, samples were visualised under an epifluorescent microscope to confirm the integrity and morphology of cells. The nucleic samples were then processed using a 10X Genomics scCNV library preparation kit following the manufacturer’s instructions.

### Genomic DNA data processing

The paired-end DNA sequencing dataset of bulk samples were first preprocessed for filtering out low quality reads, and trimming adapter bases using `fastp` [120] with default parameters. The filtered and trimmed sequences were then mapped to the mouse reference genome mm10 using `minimap2-v2.7` [121] with options `-ax sr`. Duplicated reads were marked using `MarkDuplicates` from gatk(v4.0) [122].

Single-cell DNA sequencing reads were processed using cellranger-dna cnv (v1.1.0) pipeline that demultiplexed single-cell reads and, and aligned reads to the mouse reference genome (mm10).

### Finding informative SNP markers for crossover detection

Variants in FVB/N mouse genome were downloaded from Mouse Genome Project (`FVB_NJ.mgp.v5.snps.dbSNP142.vcf`) [123], which contains the variants found in the FVB mouse genome when comparing to the reference mouse genome mm10 (C57BL/6 strain). The informative SNP markers for calling crossovers are SNPs with heterozygous genotypes in F1. To get the list of informative SNP markers, the downloaded variants were further filtered for variants with homozygous alternative genotype `(GT==1/1)`.

### Calling crossovers

Crossovers were detected by finding genotype or haplotype shifts in the BC1F1 or sperm cell genomes through modelling DNA read counts observed per sample or per cell. With DNA reads from each sample mapped and sequenced, the haplotype can be inferred by looking at the alleles carried by the DNA reads across the list of informative SNPs. To account for technical artifact (from sequencing and mapping), Hidden Markov Model based methods were implemented for crossover detection in BC1F1 samples and sperm cells with different settings (Supplementary methods).

### Hidden Markov Model

A Hidden Markov model (HMM) with binomial emission probabilities was constructed for finding crossover positions in mouse genomes while accounting for the technical artefacts including mapping errors. The observable variable in the model is the allele specific counts across the list of informative SNP sites, whereas the genotype of SNP sites is a hidden factor to be inferred. Allele read counts per sample across all informative SNP positions were obtained using `bcftools` (v.1.9) [124] with the above filtered VCF file as the region file specified via the `-R` option for each BC1F1 sample using their mapped DNA reads. Customized R scripts were implemented for applying the HMM on BC1F1 samples and inferred the sequence of hidden states against the list of SNP markers for each chromosome. Crossover detection from the single-sperm sequencing data was also by applying HMM but with different HMM configurations. Allele read counts for single cells were obtained using customized software tool *sgcocaller* [76] that summaries the allele counts across all cells and SNP positions into count matrices per sample and crossovers were called with options `–cmPmb 0.0001 –maxDP 6 –maxTotalDP 30` .

### Analysing crossover frequencies and distributions

Output files from *sgcocaller* are further parsed and analysed for crossover interval identification and crossover profile comparison. With the sequence of hidden states inferred, the crossover interval locations were called by applying functions implemented in R package *comapr* [76]. The construction of genetic distance maps and crossover distributions were also performed using *comapr*. Crossover frequencies were converted to genetic distances in centiMorgans using the Kosambi mapping function [125] via calling the `calGeneticDist` function. Permutation testing function ( `permuteDist`) was performed to test whether there is statistical difference in total or binned genetic distances between genotype groups (*Fancm ^Δ^*^2/^*^Δ^*^2^ versus *Fancm*^+/*^). (Supplementary methods)

### Meiotic chromosome spreads with immunofluorescence at prophase I

Meiotic chromosome spreads from adults testes (>8 weeks) were prepared as previously described [126]. Antibodies and dilutions used were: SYCP3 (Santa Cruz, sc-74569 at 1:100); RAD51 (EMD PC130 at 1:250); MSH4 (Abcam ab58666 at 1:200); MLH1 (BD Biosciences 51-1327GR at 1:50), MLH3 (gifted, 1:500); *γ*H2AX (Millipore at 1:500); FANCD2 (Novus Biologicals at 1:50). After immunostaining, slides were countered stained with DAPI (10 *µ*g/mL in 1xPBS) and mounted in Dako Fluorescence mounting medium. Secondary fluorescent antibodies were: goat anti-mouse Alexa Fluor 488, donkey anti-mouse Alexa Fluor 488, goat anti-mouse Alexa Fluor 594, donkey anti-rabbit Alexa Fluor 568.

Image analysis was performed using Fiji (ImageJ-win64) [127] with a macro developed by Boekhout *et al.* [128], *synapsis* [129] and manual counts. For cell scoring, the following criteria were used to classify the prophase stage: (i) leptotene: Short, discontinuous SYCP3 signals. (ii) Early zygotene: Long, discontinuous SYCP3 signals, more than 1 but fewer than 10 autosome pairs synapsing. (iii) Mid zygotene: Between 10 and 16 autosome pairs synapsed. (iv) Late zygotene: At least 17 but fewer than 19 autosome pairs synapsing, no clear XY body. (v) Early pachytene: 19 synapsed autosome pairs, clear XY body, thicker lateral element complex (as visualized by SYCP3 staining). (vi) Mid pachytene: 19 synapsed autosome pairs, clear XY body, elongated SYCP3 signal (as visualized by SYCP3 staining). (vii) Late pachytene: 19 synapsed autosome pairs, clear XY body, elongated SYCP3 signal, ending in knob-like structures (as visualized by SYCP3 staining). (viii) diplotene: At least one autosome pair starting to de-synapse.

### Meiotic metaphase I chromosome spreads

Meiotic chromosome spreads from adult testes (>8 weeks) were prepared following the methods described in [130]. For staining, slides were incubated in 10% Giemsa for 10 minutes, washed briefly with de-ionised water, air dried and mounted with DPX mounting medium.

### Micronucleus assay

Micronuclei assays were performed as described in [131].

### Fertility characterisation

*Fancm^Δ^*^2/^*^Δ^*^2^ male or female mice were assessed for fertility defects using the strategy as previously described [132]. Briefly, 8 to 12-week old mice were mated with wild-type mice for 6 months or until each pair had dropped 5 litters. Genetic combinations assessed included *Fancm*^+/+^ male x *Fancm^Δ^*^2/^*^Δ^*^2^ female, Fancm*^Δ^*^2/^*^Δ^*^2^ male x *Fancm*^+/+^ female, and *Fancm^Δ^*^2/+^ male x *Fancm^Δ^*^2/+^ female breeding pairs. Litter sizes were recorded and sex ratios were monitored.

Testes were harvested from adult mice and fixed in Bouin’s solution (Amber Scientific) for 5 hours at room temperature and alcohol processed for histopathology into paraffin wax and sectioned using standard methods. Sectioned dewaxed testis sections were stained using periodic acid-Schiff’s reagent and haemotoxylin reagents. All slides were dehydrated and mounted under a coverslip with DPX (Sigma-Aldrich, USA).

Testis daily sperm production (DSP), was determined as previously described [133]. Briefly harvested testes were weighed, snap-frozen and stored at -80°C. After thawing, testes were homogenized by sonication in DSP buffer (0.15 M NaCl, 0.01% NaN3, 0.05% Triton X-100) to lyse all cells except the condensed spermatid nuclei. The number of elongated spermatids per testis were determined used a haemocytometer and DSP estimated by dividing the total elongated spermatid number per testis by 4.84, which corresponds to the estimated number of days that an elongated spermatid remains within the testis [134].

Seminiferous tubule analysis was performed on Bouin’s fixed and periodic acid-Schiff’s reagent and haemaotoxylin stained testis sections using FIJI software. Image acquisition was performed with a Leica Thunder microscope and LASX software.

### Computer-assisted sperm analysis (CASA)

Analysis of sperm motility was conducted as previously described [135]. Briefly, cauda epididymal spermatozoa were collected via backflushing of 10-12 week old mice. Sperm were transferred into modified Tyrode’s 6 medium and loaded into 80 *µ*m deep CASA slides. Movement parameters were measured using the MouseTraxx CASA system (Hamilton-Thorne, USA). Rapid movement was classified as velocity over 35 *µ*m/s, medium motility was classified as 10–35 µm/s, slow motility was classified as *<*10 µm/s and static was 0 µm/s. At least 1000 sperm were measured per animal with a minimum of four mice per genotype and age.

### Direct follicle counts

Ovarian follicles were quantified as previously described [136]. Briefly, ovaries were harvested and fixed in either 10% (v/v) neutral buffered formalin solution (#ANBFC, Australian Biostain) or Bouin’s solution (#HT10132, SigmaAldrich) for 24 hours. Tissue was embedded in paraffin and was exhaustively serially sectioned at 5 µm with a MicroTec Cut 4060 paraffin microtome and every ninth section was collected and stained with periodic acid-Schiff’s reagent and haematoxylin. Slides were scanned at 20x using an Aperio Digital Pathology Slide Scanner (Leica Biosystems). The Aperio Imagescope program was used to quantify every primordial, primary, secondary, antral and atretic follicle based on morphology [137] and for which an oocyte nucleus was clearly visible, to obtain raw counts of oocytes sampled. The total follicle number was determined by multiplying the raw counts by 9 to correct for the sections not counted. Direct follicle count data was analysed using Graphpad Prism 8 (Version 8.0.2). These data are presented as mean *±* SEM and p-values for each graph are specified following unpaired t-tests.

### Oocyte collection

For superovulation, adult female mice were injected subcutaneously with 5 IU equine chorionic gonadotropin, followed 44 to 48 h later with 5 IU human chorionic gonadotropin. Oocytes were harvested from oviducts 12 to 14 h later.

### Germ cell apoptosis analysis

Germ cell apoptosis was evaluated by immunostaining of cleaved caspase-3 (Cell signalling #9664, 1:100) and -7 (Cell signalling #9491, 1:500). 5 *µ*m testis sections were dewaxed and antigen retrieval was performed using citrate buffer pH 6.0. Sections were blocked for endogenous peroxidase with 3% H_2_O_2_ for 5 min and then washed with 1xPBS twice. Sections were then blocked with Cas-Block^TM^ (ThermoFisher #008120) for a minimum of 30 minutes at room temperature. Sections were then incubated in primary antibodies in Dako antibody diluent) overnight at 4*^◦^*C. The next day, sections were washed three times for 5 min. Secondary anti-rabbit Dako Envision Dual Link-HRP (#K4063) was incubated on the sections for 30 min at room temperature, then washed three times for 5 min with 1xPBS. DAB Dako liquid chromagen was applied for 1 min, then removed and washed with water for 5 min. Sections were counterstained with Mayer’s Haemotoxlyin for 4 min and washed briefly with water. Scott’s Tap Water was applied to the sections for 1 min then washed briefly with water. Sections were then dehydrated, cleared and mounted with DPX mounting medium.

Sections were then scanned and the number of cleaved caspase-positive cells in a minimum of 90 randomly selected seminiferous tubules per mouse were counted (n = 3 per genotype).

Statistical analysis of germ cell apoptosis data was then conducted in R version 3.5.1 [138]. Generalised linear mixed (GLM) models are used to compare the number of caspase-positive cells per tubule across genotypes, with animal ID included as a random effect to account for repeated measures per individual. For each model, Akaike information criteria (AIC) estimates are used to select the most appropriate error distribution and link function (i.e. Poisson, negative binomial, zero-inflated Poisson, zero-inflated negative binomial) using the glmer function (lme4 package; [139] ) and glmmTMB function (glmmTMB, package; [140]). For all models, a zero-inflated negative binomial distribution (fitted with glmmTMB, using the ziformula argument) was selected as the most appropriate error distribution and link function (i.e. had the lowest AIC score).

### MMC-induced chromosomal breakage analysis

MMC-induced chromosomal breakage tests were assessed using primary splenic B lymphocytes from *Fancm* mice using methods from [141, 142]. After culturing B cells in the presence of LPS for 24 hours, mitomycin C was added at varying concentrations (10, 50 and 100 ng/mL). After 72 hours, cells were given fresh media containing colcemid (Karyomax, 0.2 *µ*g/mL) for two hours. Following colcemid treatment, cells were harvested, washed and resuspended in 1x PBS. Pre-warmed swelling buffer (75 mM KCl) was added by dropwise addition while gently vortexing. Cells were incubated at 37*^◦^*C for 10 mins and a couple of drops of chilled fixation buffer (1 methanol : 1 glacial acetic acid) was added. Cells were spun down and supernatant removed. Cells were resuspended in chilled fixative buffer with constant agitation three times before dropping spreading onto slides.

### Delivery of ibuprofen in drinking water

Ibuprofen treatment was performed following methods described in McNairn *et al.* [7]. *Fancm*^+/^*^Δ^*^2^ males were mated to *Fancm*^+/^*^Δ^*^2^ females. The parents and pups were provided with water bottles containing ibuprofen (children’s Advil), 5 mL (100 mg) in 250 mL. They were allowed to drink ad libitum (50-80 mg per kg per day) at all stages of development. Newborn mice were genotyped for the *Fancm* allele. Gonads and blood were collected from 8-week old pups for analysis.

## Acknowledgments

We are grateful to Chloé Girard and Andrew Lloyd and Raphaël Mercier for helpful comments on the manuscript. Florencia Pratto, Kevin Brick and Phil Jordan provided assistance and helpful feedback with the crossover density predictive modelling. The surface spreading technique was generously taught by Rocío Gómez. WC and DM receive Fellowships and funding related to this work from the Australian National Health and Medical Research Council (GNT1129757, GNT1112681, GNT1185387). RL and VT are recipients of a Research Training Program Scholarship from the Australian Commonwealth Government and the University of Melbourne and SVI Foundation Top-Up Scholarship from St Vincent’s Institute. RL receives a Xing Lei PhD Top-up Scholarship in Mathematics and Statistics. VT receives a St Vincent’s Institute Top-Up scholarship. LM received a fellowship from the Lorenzo and Pamela Galli Research Trust. The anti-MLH3 antibody was a gift from Corinne Grey, Bernard de Massy and Valérie Borde. The *Sting* mouse line was a gift from Benjamin Kile. Casey Anttila, Peter Hickey, Stephen Wilcox from the WEHI Cellular Genomics Projects and Sequencing Teams assisted with scRNA-seq experiments. The authors would like to acknowledge the technical support of the Monash Animal Research Platform and Monash Histology Platform, Monash University. BioRender was used for illustrations.

## Competing interests

The authors declare no competing interests.

## Supporting information

**S1 Fig.**
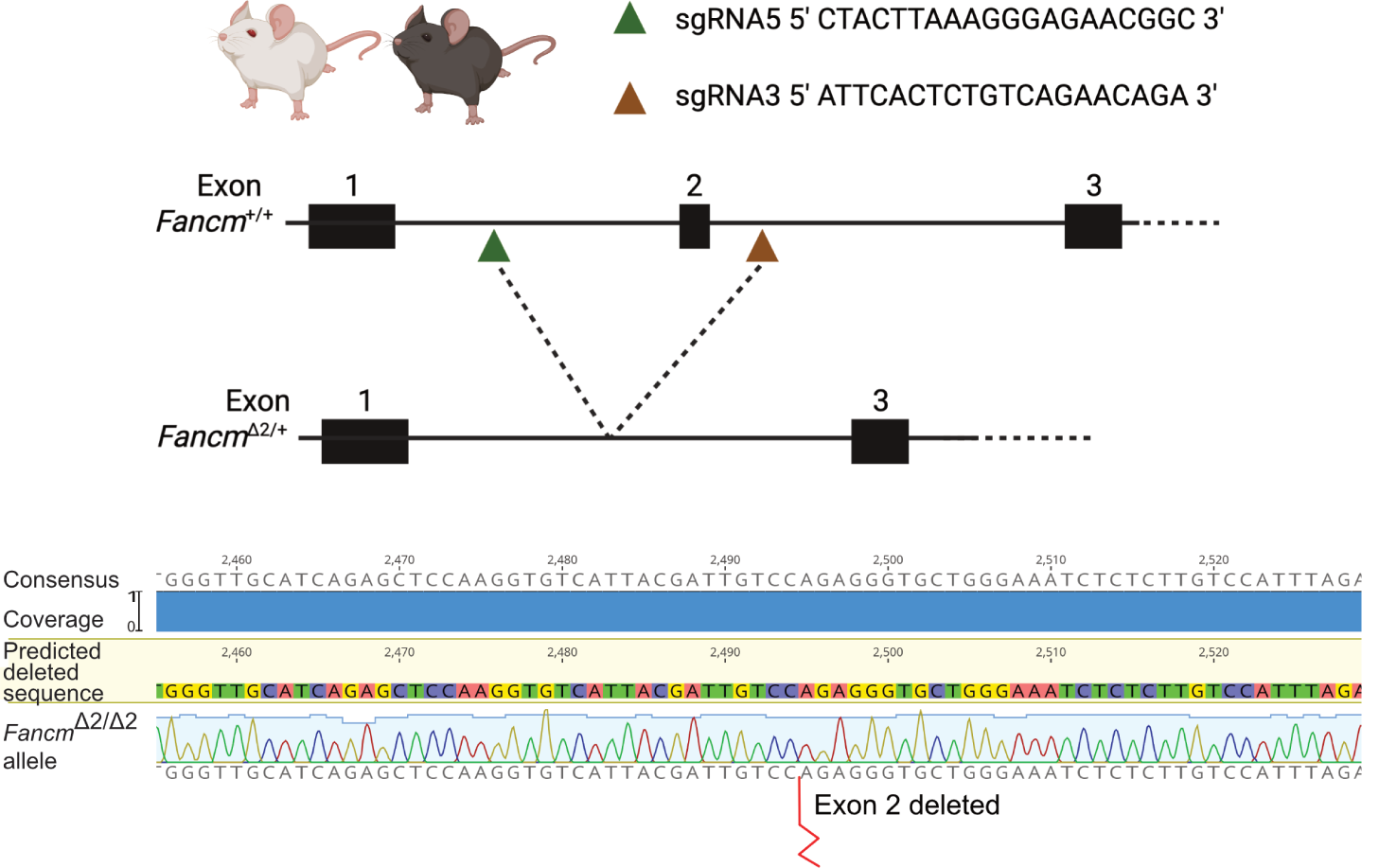
Transgenic mouse generation. *Fancm* mouse knockouts were made in two strains, C57BL/6J and FVB/N, with the same two guide RNAs, which resulted in the targeted deletion of exon 2.

**S2 Fig.**
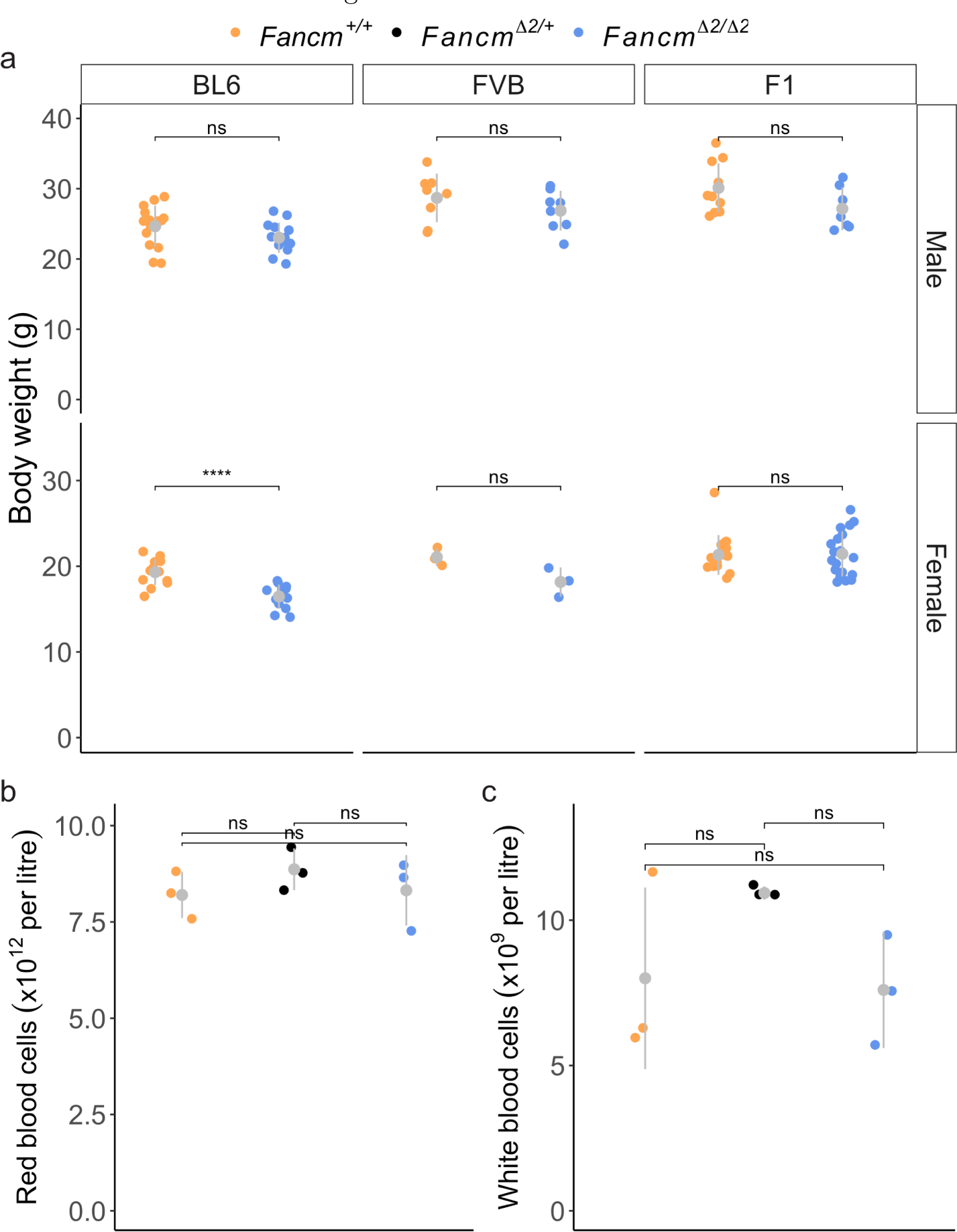
Body weight and blood parameters are normal in *Fancm* ^*Δ*2/*Δ*2^ mice. a) Body weights of 6-14 week old *Fancm*-deficient mice and control litter mates. (**** indicates p-value *≤* 0.0001 (unpaired t-test). b) Red blood counts from B6.*Fancm* mice and their control siblings. c) White counts from B6.*Fancm* mice and their control siblings.

**S3 Fig.**
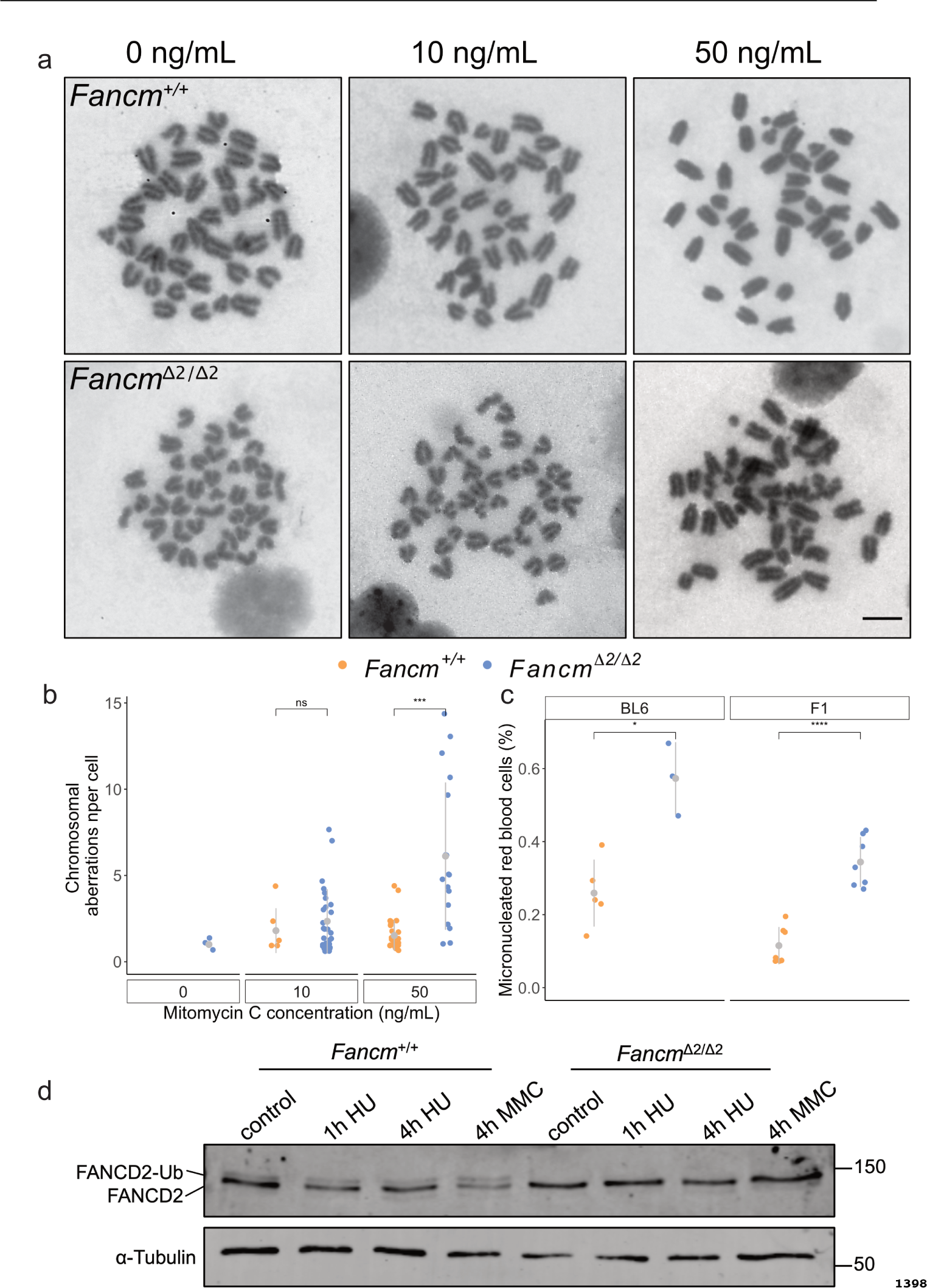
Genomic instability in *Fancm* -deficient mice. a) Chromosome breakage tests shows a dose-dependent response to mitomycin C (0, 10 and 50 ng/mL) in the absence of *Fancm*. Scale bar is 5 *µ*m. b) Quantification of chromosomal aberrations per in *Fancm*^+/+^ and *Fancm^Δ^*^2/^*^Δ^*^2^ cells. c) Increased micronuclei in mature red blood cells reveals an increase in genomic instability in *Fancm* mutants. d) FANCD2 mono-ubiquitination in splenic B cells measured by western blot. An 8 kDa shift occurs when FANCD2 is mono-ubiquitinated. This can occur under basal conditions and treatment with hydroxyurea (10 mM) which causes replication fork stalling, or the DNA crosslinker mitomycin C (50 *µ*g/*µ*L). FANCD2 mono-ubiquitination is not detected in the absence of *Fancm* under any of these conditions. Error bars indicate mean *±* s.d., *** indicates p *≤* 0.001, unpaired t-test.

**S4 Fig.**
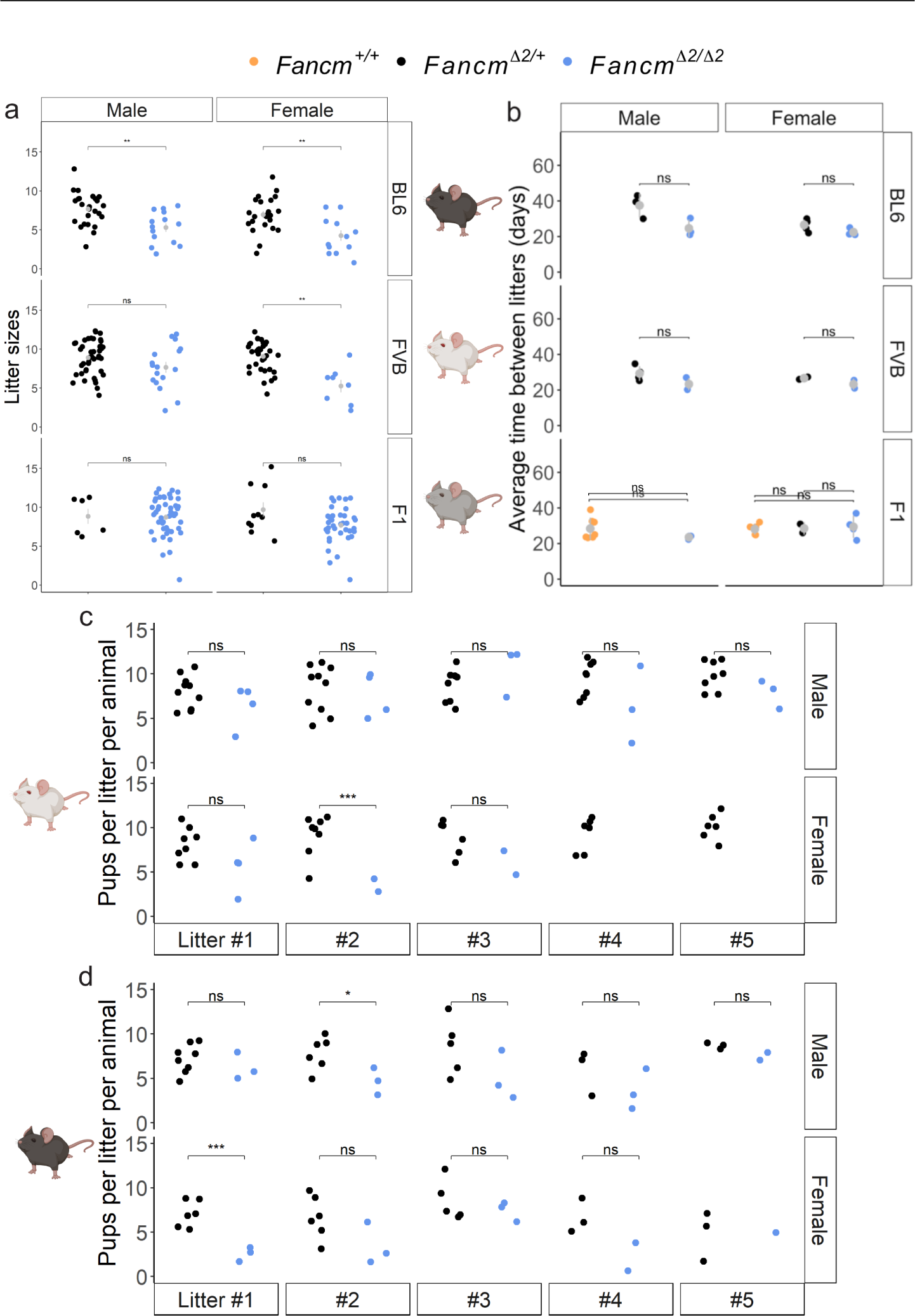
Litter sizes from *Fancm* -deficient mice and litter mate controls. a) Average litter sizes over five litters per mouse in C57BL/6J, FVB/N and F1 strains. b) Average time between litters in days in C57BL/6J, FVB/N and F1 mice. The average pups per litter was also considered in c) FVB/N and b) C57BL/6J mice. Mice were allowed to breed for five litters or 6 months. For the C57BL/6J and FVB/N data, their mate of the opposite sex had a wild-type *Fancm* allele. In the case of F1 mice, their mate of the opposite sex had either a wild-type or heterozygous *Fancm* allele. Data shown are mean *±* s.d. * indicates p *≤* 0.05, ** indicates p *≤* 0.01, *** indicates p *≤* 0.001, unpaired t-test.

**S5 Fig.**
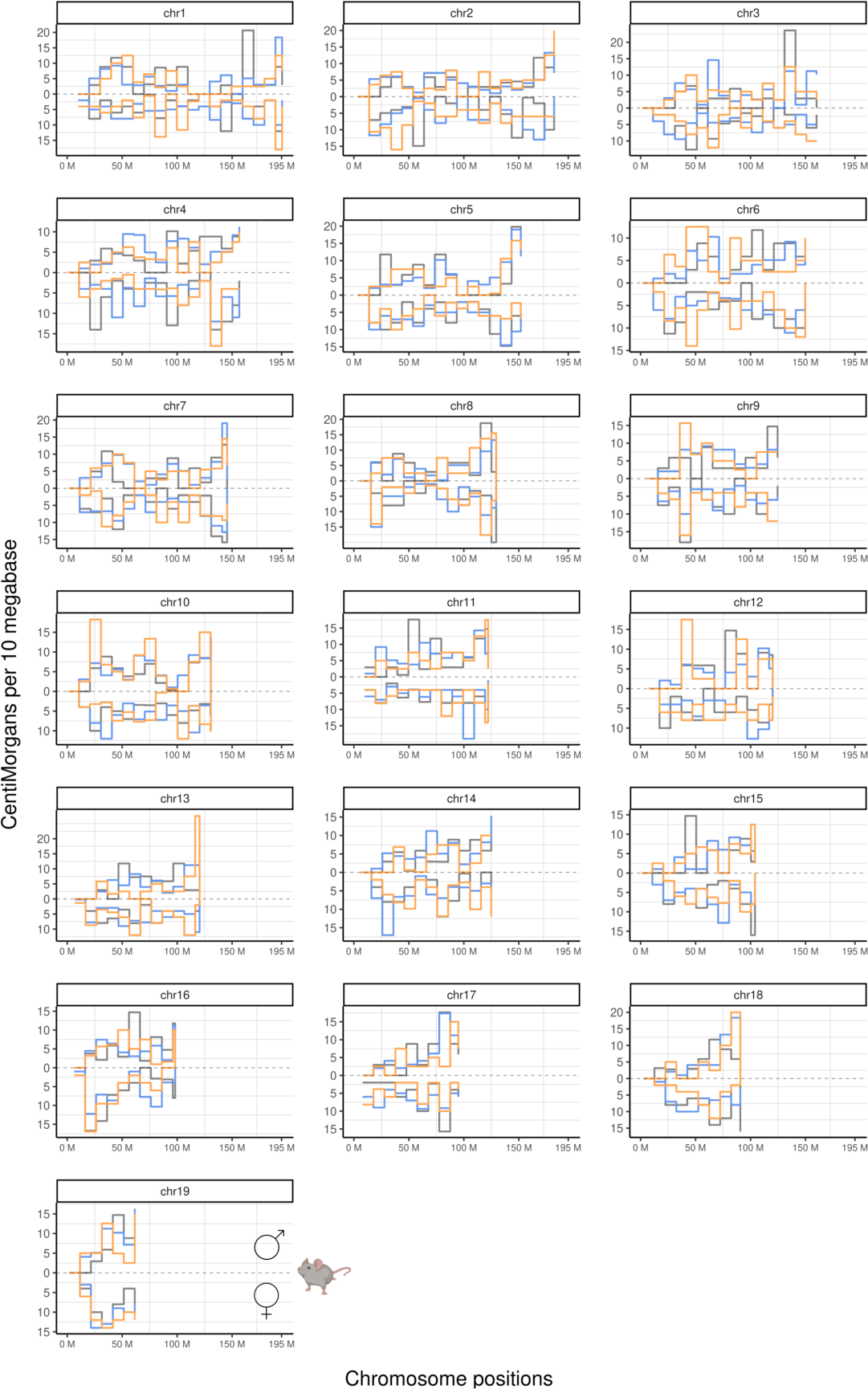
Male and female genome-wide genetic distances. Male and female crossover frequencies detected using bulk sequencing of pedigree samples. Genetic distances in each bin between *Fancm*^+/+^ (orange), *Fancm^Δ^*^2/+^ (black) and *Fancm^Δ^*^2/^*^Δ^*^2^ (blue) were tested, and there were no significant differences (permutation testing, B = 1,000).

**S6 Fig.**
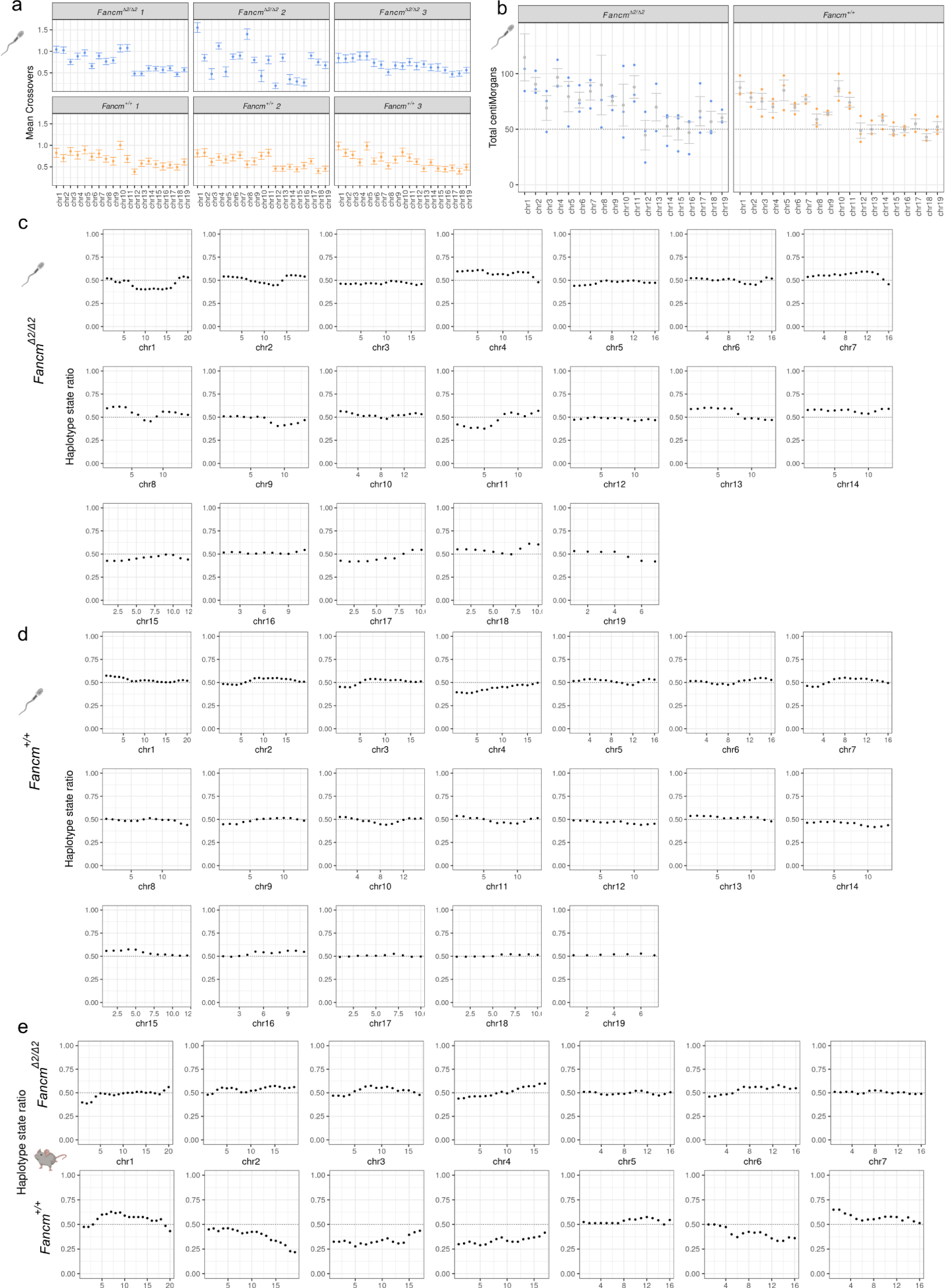
Crossover detection with single sperm sequencing. Droplet-based single sperm sequencing produced datasets that could detect crossovers. a) Mean crossovers per chromosome of sperm sequenced from F1 individuals. b) Total centiMorgans per chromosome from observed crossover rates per individual. Number of cells per individual: *Fancm^Δ^*^2/^*^Δ^*^2^: 114, 40, 64. *Fancm*^+/+^ 57, 70, 63. c) Marker segregation ratio in all gametes from *Fancm^Δ^*^2/^*^Δ^*^2^ F1 individuals. Ratios of counts of cells with the two haplotype were calculated in chromosome bins (of size 10 mega bases). d) Marker segregation ratio in all gametes from *Fancm*^+/+^ F1 individuals. Hypothesis testing using binomial test was perform to test if the marker segregation ratio is different from 0.5, no significant differences were observed in any chromosomes. e) Same observation in Bulk BC1F1 pups that no marker segregation distortions was found (only showing selected chromosomes).

**S7 Fig.**
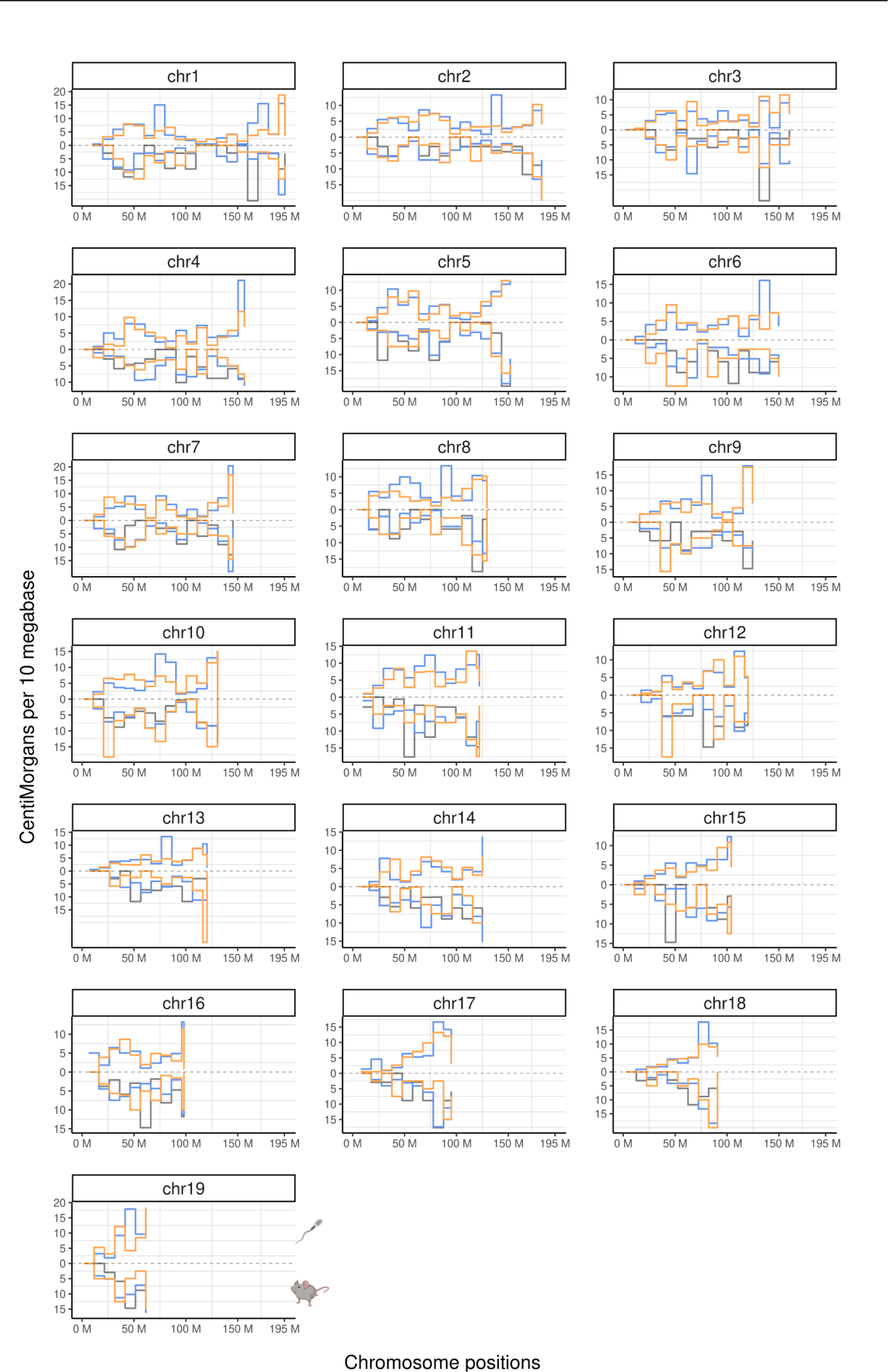
F1 single-sperm sequencing and BC1F1 bulk sequencing produce comparable crossover distributions. Male recombination maps of all 19 mouse autosomes are presented in separate plots of *Fancm*^+/+^ (orange), *Fancm^Δ^*^2/+^ (black) and *Fancm^Δ^*^2/^*^Δ^*^2^ (blue). The genetic distances from sperm-sequencing data from F1 mice is flipped to the negative scale. Bin sizes are 10 megabases.

**S8 Fig.**
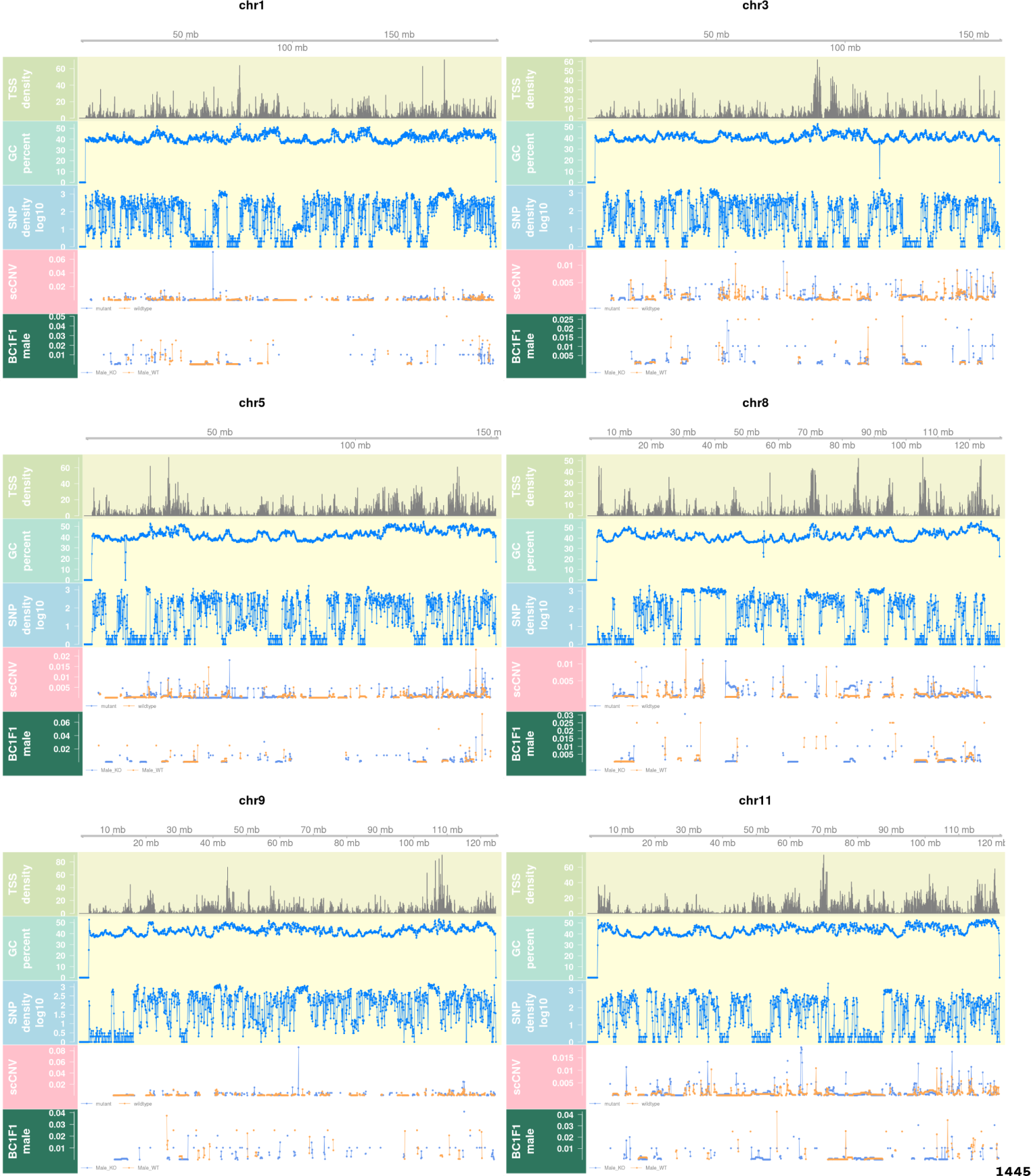
The effect of chromosomal features on crossover frequencies. Visual representation of the effect of SNP density and other chromosomal features on crossover frequency (Size of 100kb physical base pair bins) from selected chromosomes. The bottom two rows of each chromosome track represent crossover densities for the single-gamete sequencing (“scCNV”) and pedigree-based (“BC1F1 male”) methods. Orange = *Fancm*^+/+^ and blue = *Fancm^Δ^*^2/^*^Δ^*^2^.

**S9 Fig.**
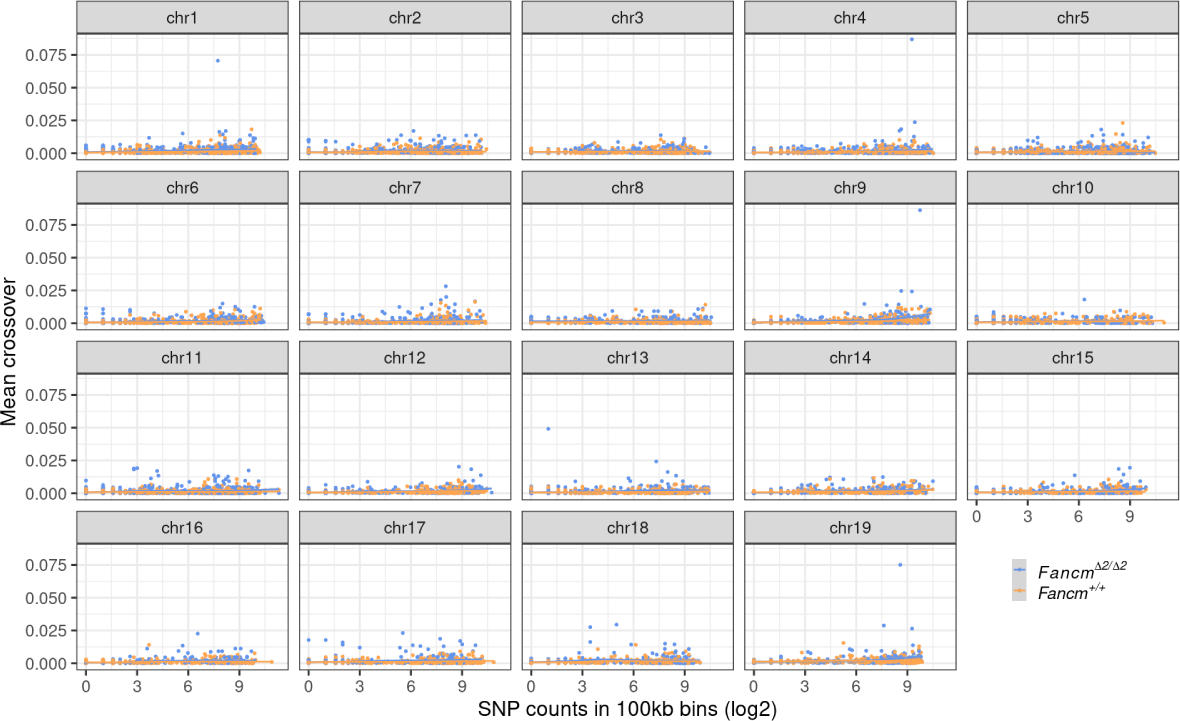
Assessment of the relationship between SNP density and crossov frequency. Visual representation of the effect of SNP density on crossover frequency using a physical bin size of 100kb).

**S10 Fig.**
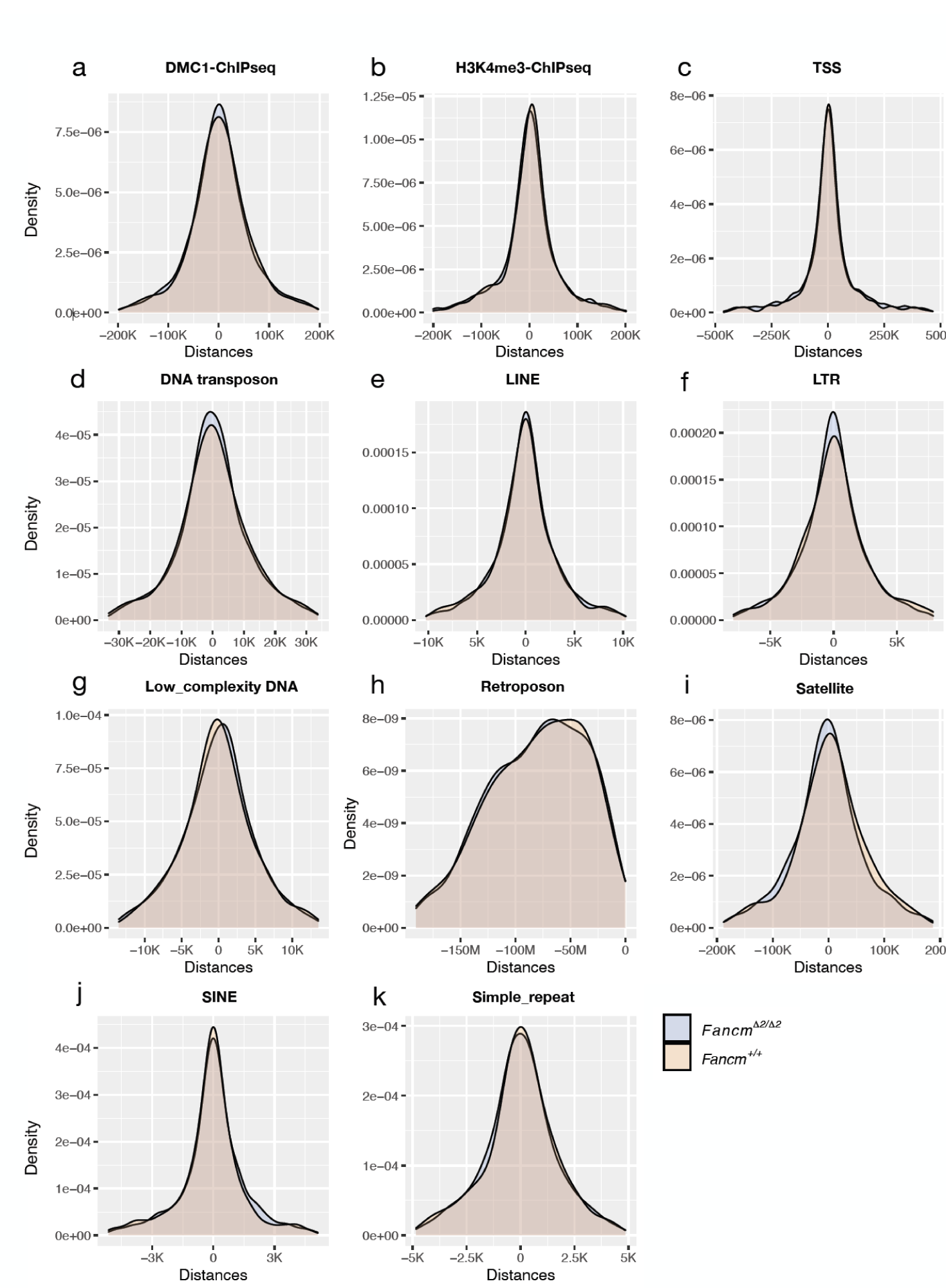
The correlation of meiotic DSBs and genomic features on crossover frequency. Meiotic crossover frequency was compared to DSBs and genomic features to determine if the extra crossovers in *Fancm*-deficient mice display typical patterns of distribution with respect to these variables. a) DMC1ChIPseq data from [143], which is a proxy for DSBs, b) H3K4me3-ChIPseq data from [143], which is commonly associated with accessible chromatin and crossover occurence, c) transcription start sites, d) DNA transposons, e) long interspersed nuclear elements, f) long terminal repeat retrotransposons, g) low complexity repeats are AT-rich or GC-rich regions, h) retrotransposons, i) satellite repeats, j) short interspersed nuclear elements, k) simple repeats are microsatellite repeats that have interspersed characters.

**S11 Fig.**
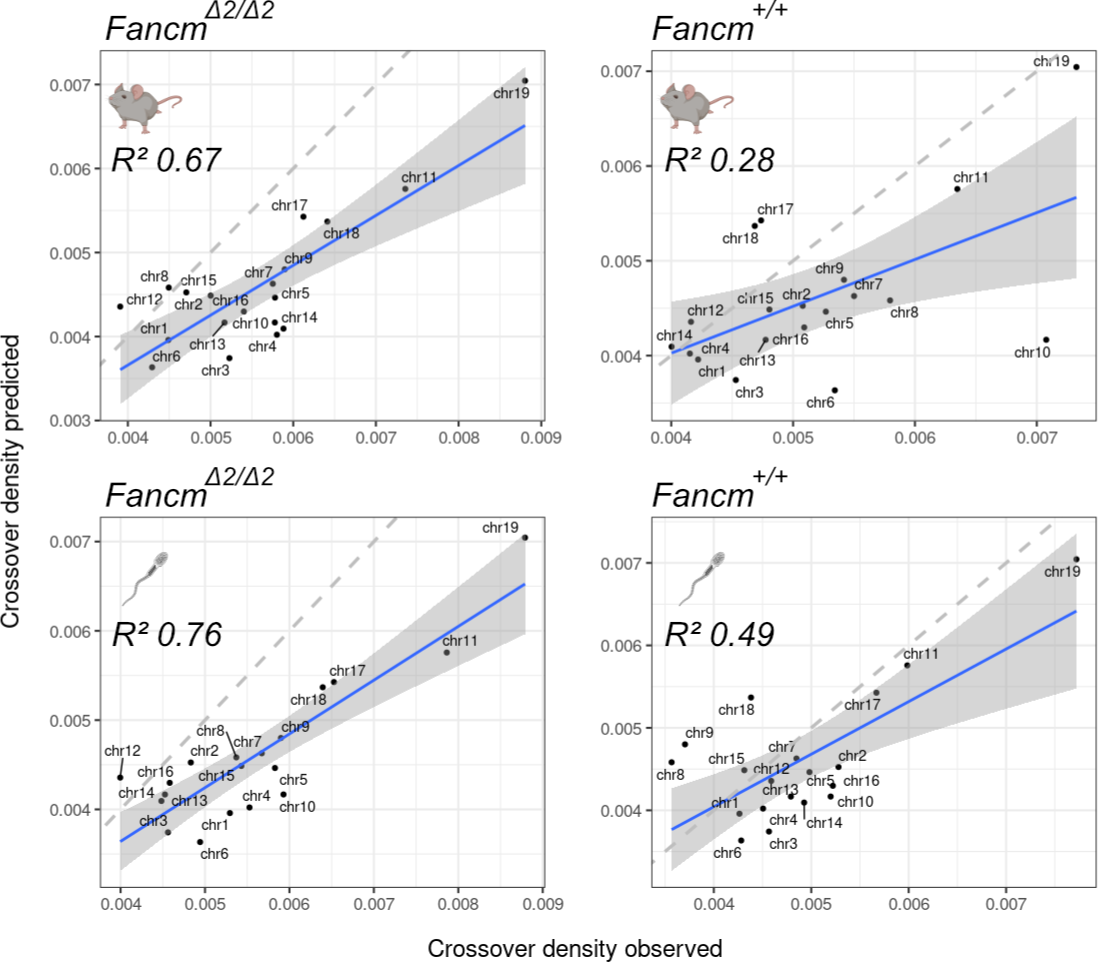
Experimental and predicted regression modelling of crossover densities. The model from [29] was used to predict crossover density for the 19 autosomes. Linear regression was fitted to compare the predicted crossover densities with observed crossover densities from each sample and data group. Dashed lines indicate the diagonal lines with slope 1 with adjusted R squares printed generated using lm() from R [138]. Grey area is the standard error of the regression.

**S12 Fig.**
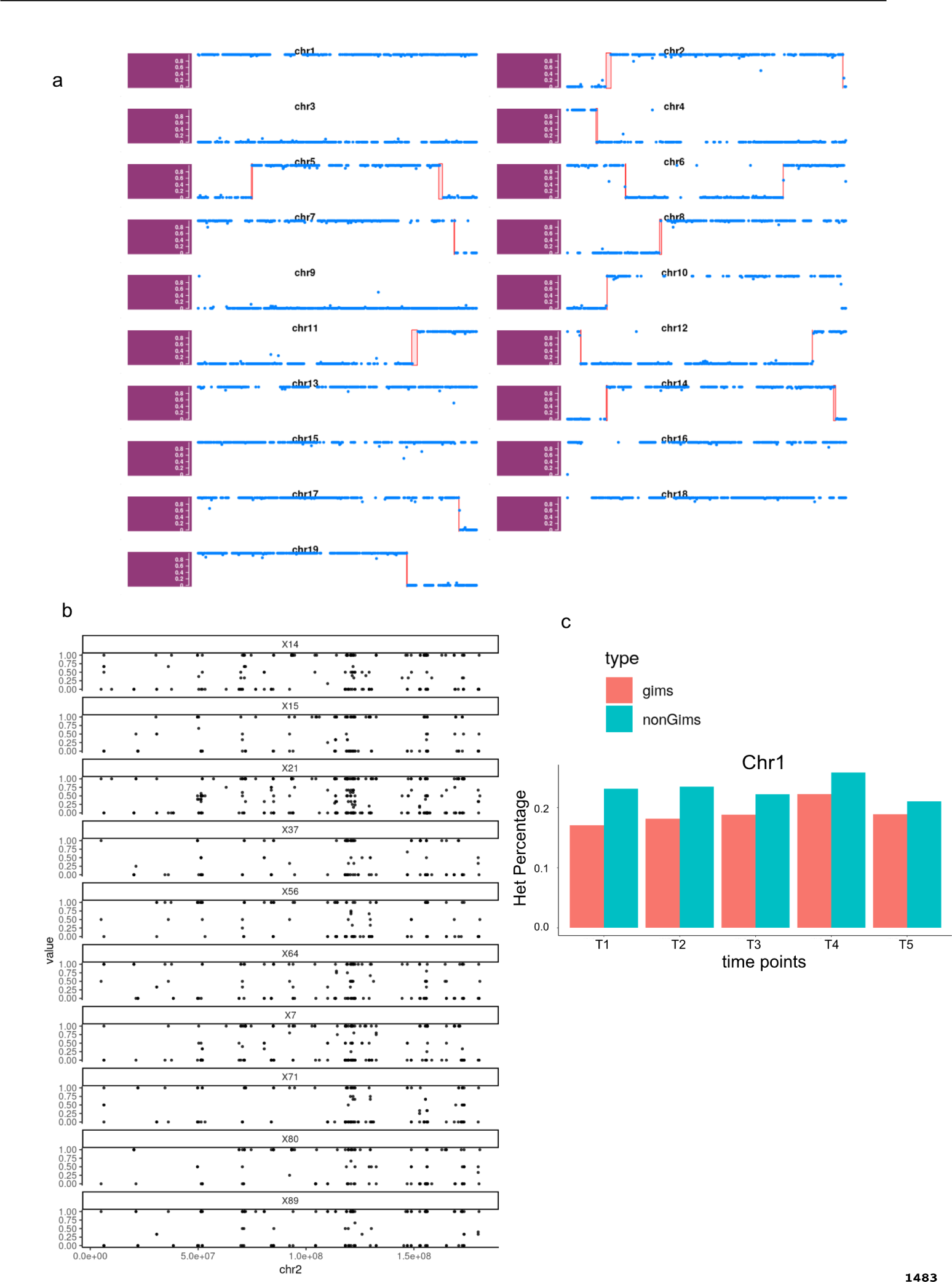
Mendelian inheritance of DNA, but not RNA, in haploid sperm. a) Single sperm DNA-sequencing methods produce data that is sufficient for crossover detection, b) whereas scRNA-sequencing methods do not. c) However, selected transcripts (GIMs from [80]) are more representative of the haploid nuclear allelic content, but still show significant transcript sharing, which is reflected by detection of heterozygosity. 100 cells from each time points are selected and the SNPs were grouped by whether falling in GIM or non GIMs regions. The percentage of the SNPs were called with heterogzygous (alternative allele frequencies were between 0 (C57BL/6J) to 1 (FVB/N) were plotted).

**S13 Fig.**
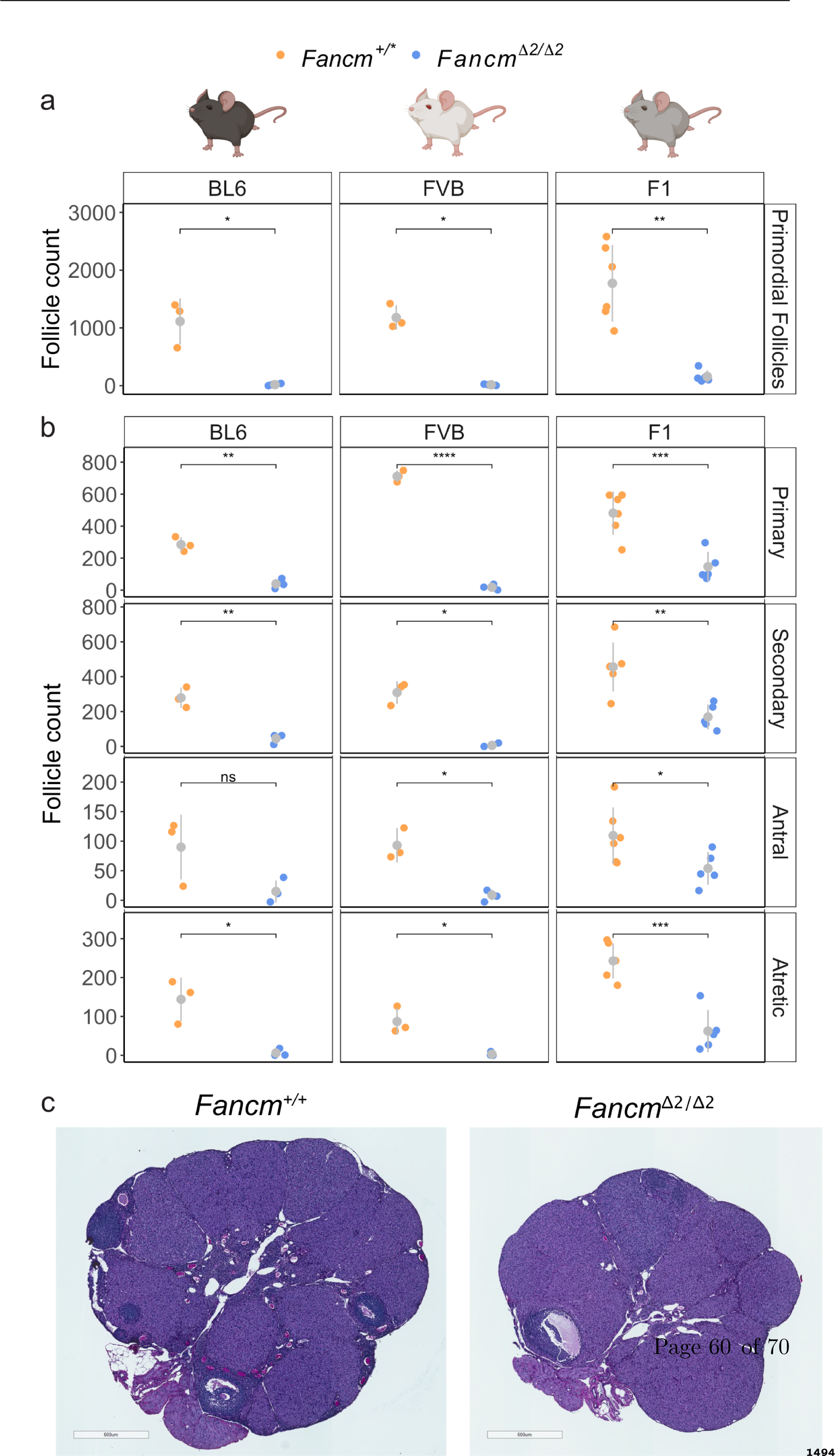
Folliculogenesis is perturbed *Fancm* -deficient mice. Follicle numbers in 3 month old *Fancm*^+/*^ and *Fancm^Δ^*^2/^*^Δ^*^2^ in C57BL/6J, FVB/N and F1 strains. a) Primordial follicle number are significantly reduced in FVB.*Fancm^Δ^* B6.*Fancm^Δ^*^2/^*^Δ^*^2^ and F1.*Fancm^Δ^*^2/^*^Δ^*^2^ ovaries. b) Primary, secondary and atretic follicle numbers are significantly lower in FVB.*Fancm^Δ^*^2/^*^Δ^*^2^, B6.*Fancm^Δ^*^2/^*^Δ^*^2^ F1.*Fancm^Δ^*^2/^*^Δ^*^2^ ovaries. c) Representative images of middle sections of FVB.*Fanc* and FVB.*Fancm^Δ^*^2/^*^Δ^*^2^ ovaries. Scale is 600 *µ*m. *Fancm*^+/*^ indicates *Fancm*^+/+^ or *Fancm*^+/^*^Δ^*^2^. Data shown are mean *±* s.d. * indicates p *≤* 0.05, ** indicates p *≤* 0.01, *** indicates p *≤* 0.001.

**S14 Fig.**
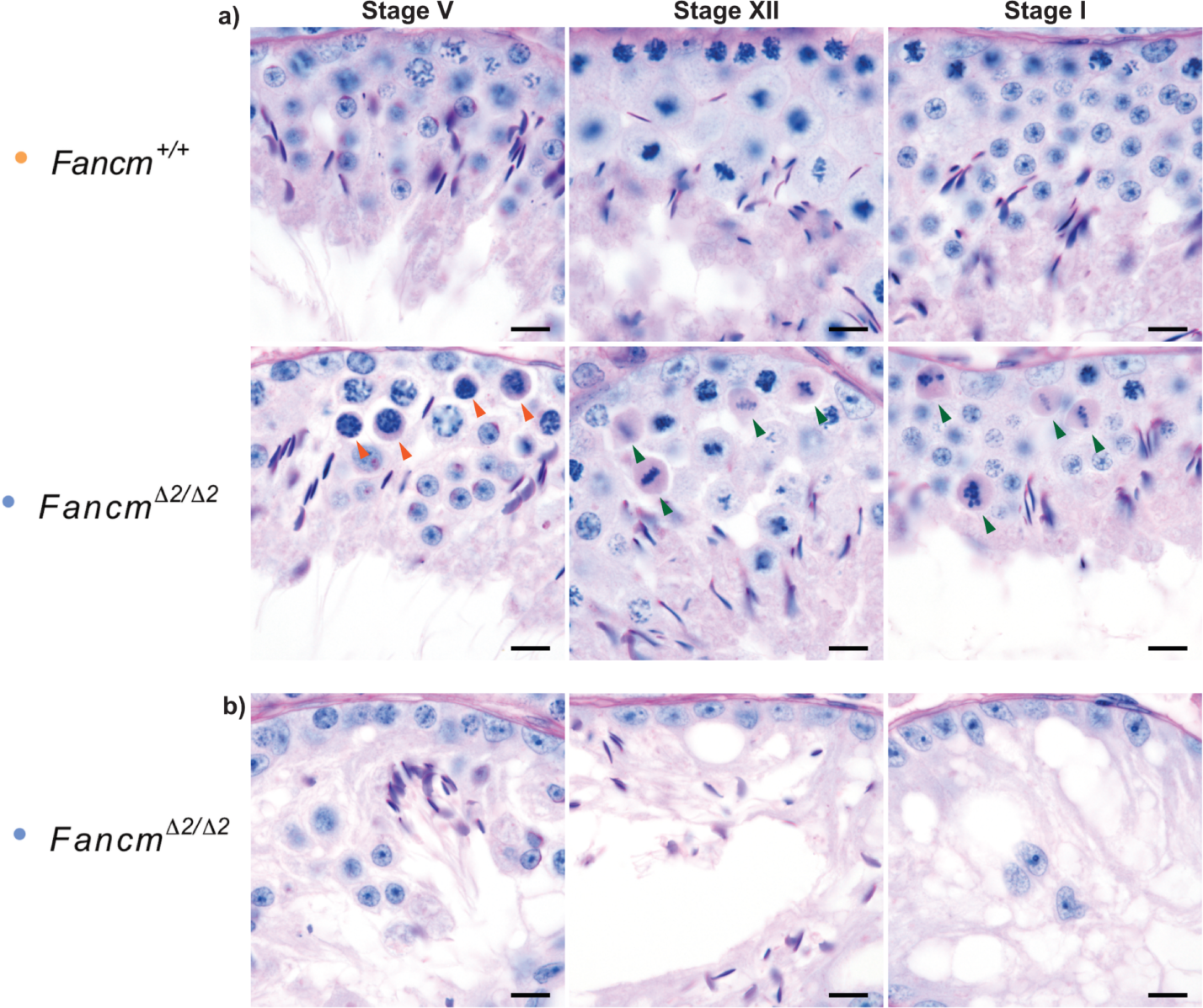
Representative images of histological defects observed in B6.*Fancm*^*Δ2/Δ2*^ seminiferous tubules. a) Orange arrows indicate PAS-positive pachytene spermatocytes, green arrows indicate PAS-positive spermatocytes in stage XII and I. b) Representative images that depict tubules where spermatocytes appear to be the first population to be lost. On the left panel spermtogonia, Sertoli cells, spermatids (round and elongated) were observed but no spermatocytes. The middle and far right panels show progression to a Sertoli cell-only phenotype. Scale bar = 10 microns.

**S15 Fig.**
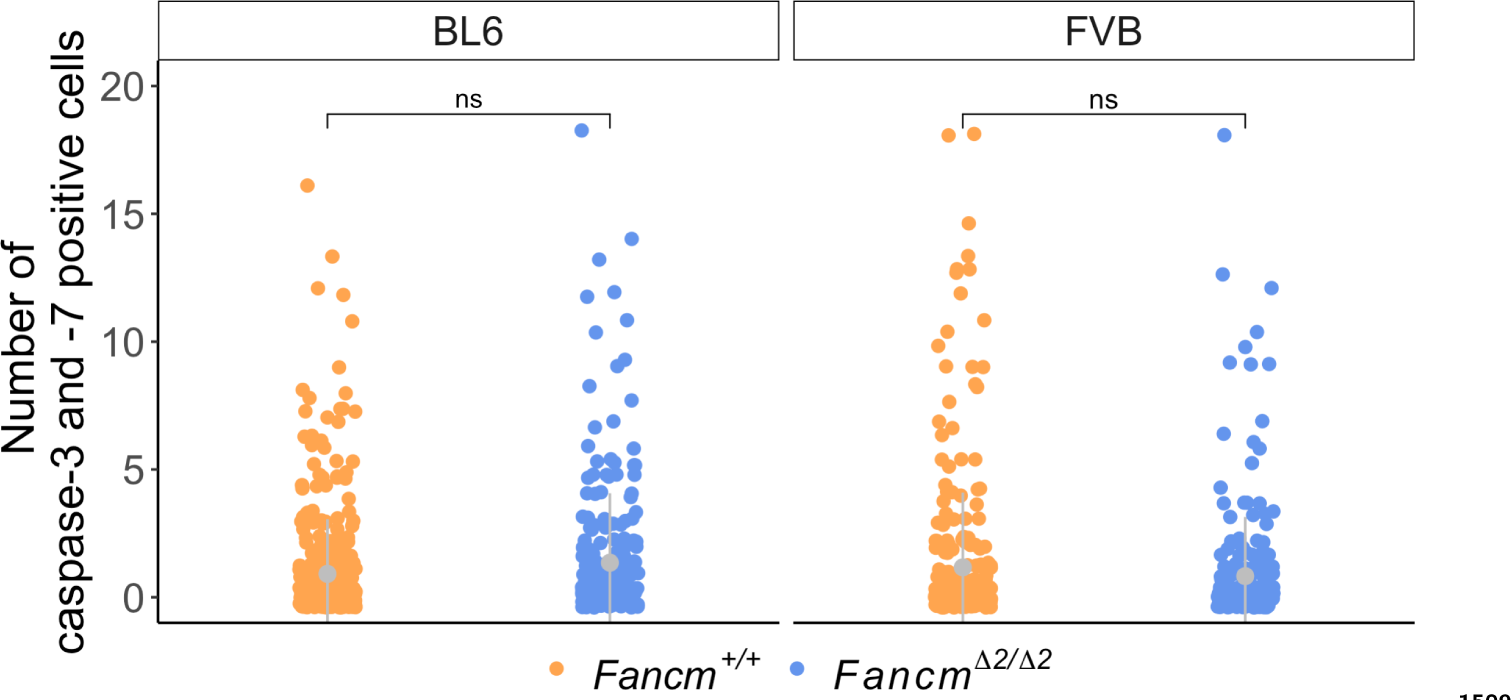
Germ cell apoptosis analysis in *Fancm* -deficient seminiferous tubules. Caspase-3 and -7 positive cells *Fancm^Δ^*^2/^*^Δ^*^2^ in BL6 and FVB seminiferous tubules are not significantly different from *Fancm*^+/+^. Generalised linear mixed (GLM) models used as to compare the number of caspase-positive cells per tubule between genotypes.

**S16 Fig.**
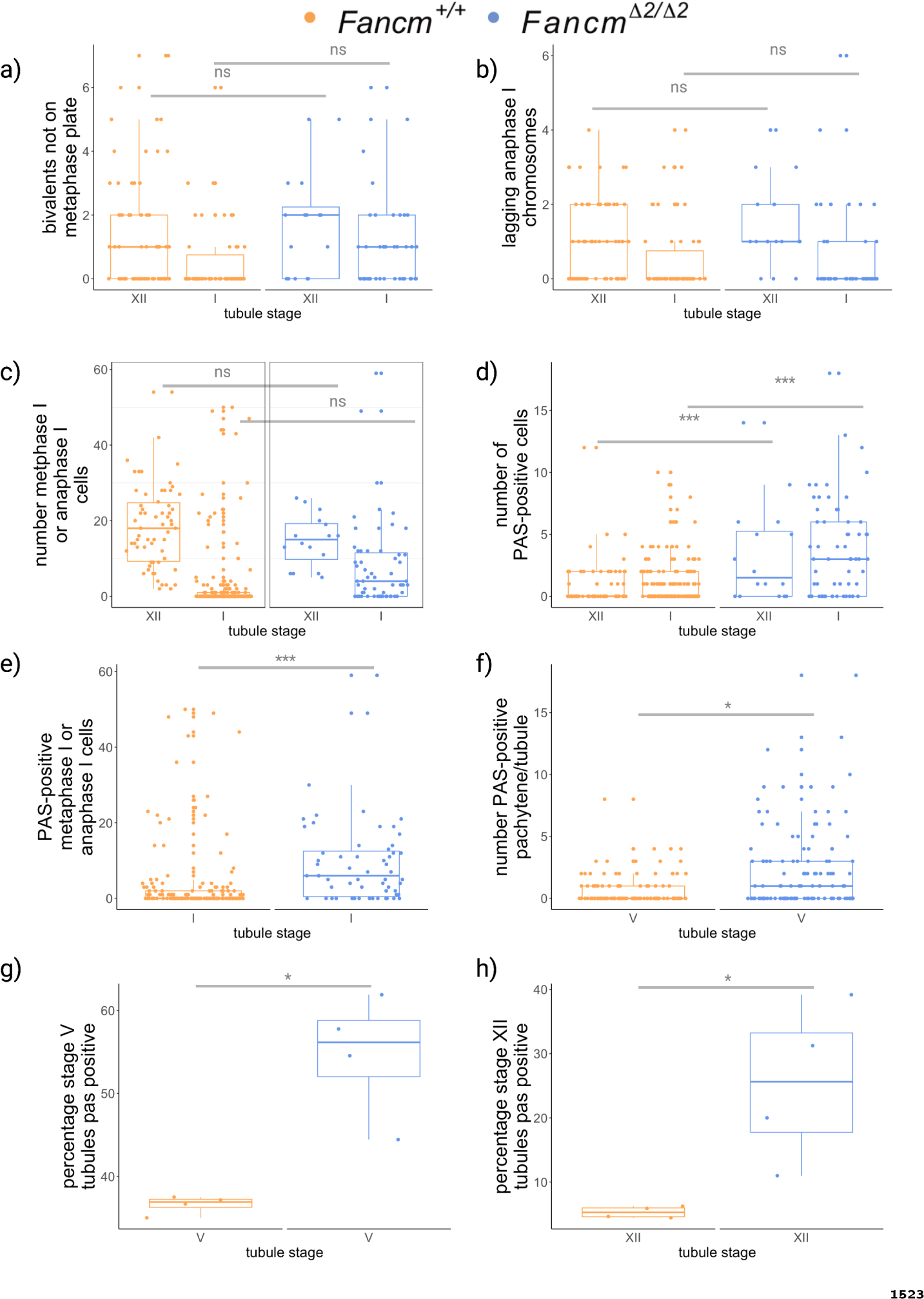
Increased histological defects *Fancm* seminiferous tubules. PAS-stained histological sections of seminiferous tubules were used for quantification of abnormalities. a) Bivalents align on the metaphase plate and b) segregate at anaphase I comparably in B6.*Fancm^Δ^*^2/^*^Δ^*^2^ and B6.*Fancm*^+/+^. c) The number cells in metaphase I and anaphase I in stage XII and stage I seminiferous tubules was comparable between genotypes. d) An increase in PASpositive or pyknotic cells was observed in both stage XII and stage I tubules in B6.*Fancm^Δ^*^2/^*^Δ^*^2^. e) PAS-positive pachytene and f) metaphase I/anaphase I were increased in B6.*Fancm^Δ^*^2/^*^Δ^*^2^. g) The percentage of stage V tubules containing PAS-positive pachytene spermatocytes. h) The percentage of metaphase I-anaphase I cells spermatocytes that were PAS-Positive in Stage XII. * indicates p *≤* 0.05, ** indicates p *≤* 0.01, *** indicates p *≤* 0.001.

**S17 Fig.**
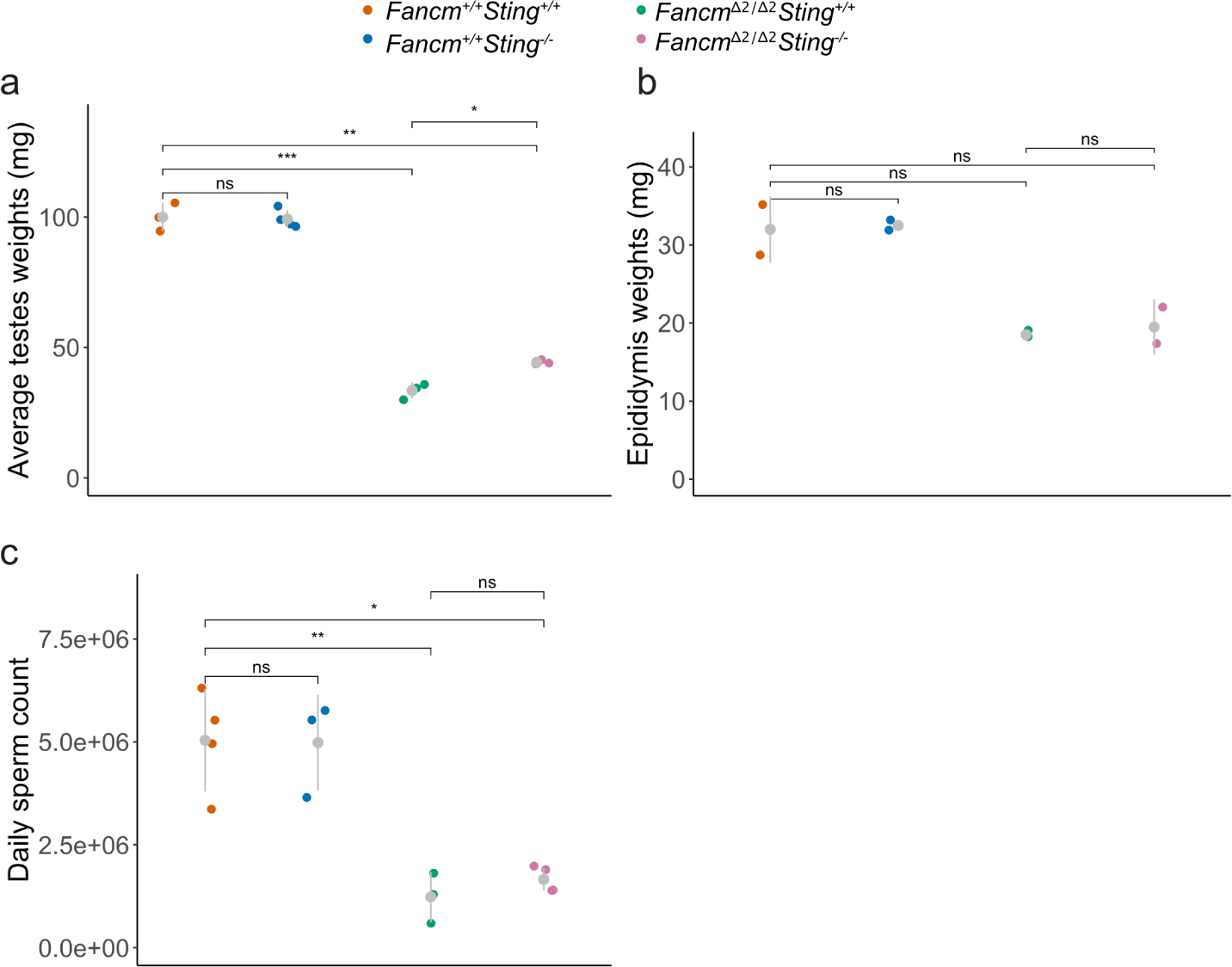
Sting depletion partially rescues gonad weights *Fancm* -deficient mice. a) Testicular weight exhibited a subtle partial but significant rescue in *Fancm^Δ^*^2/^*^Δ^*^2^*Sting* ^-/-^ compared to *Fancm^Δ^*^2/^*^Δ^*^2^. b) epididymal weight and c) daily sperm production were not significantly different between *Fancm^Δ^*^2/^*^Δ^*^2^*Sting* and *Fancm^Δ^*^2/^*^Δ^*^2^*Sting* ^-/-^. Due to limited epididymis sample numbers, there is low statistical power to detect difference. * indicates p *≤* 0.05, ** indicates p *≤* 0.01, *** indicates p *≤* 0.001 and *** indicates p *≤* 0.0001 (one-sided t-test with multiple testing correction “fdr”).

**S18 Fig.**
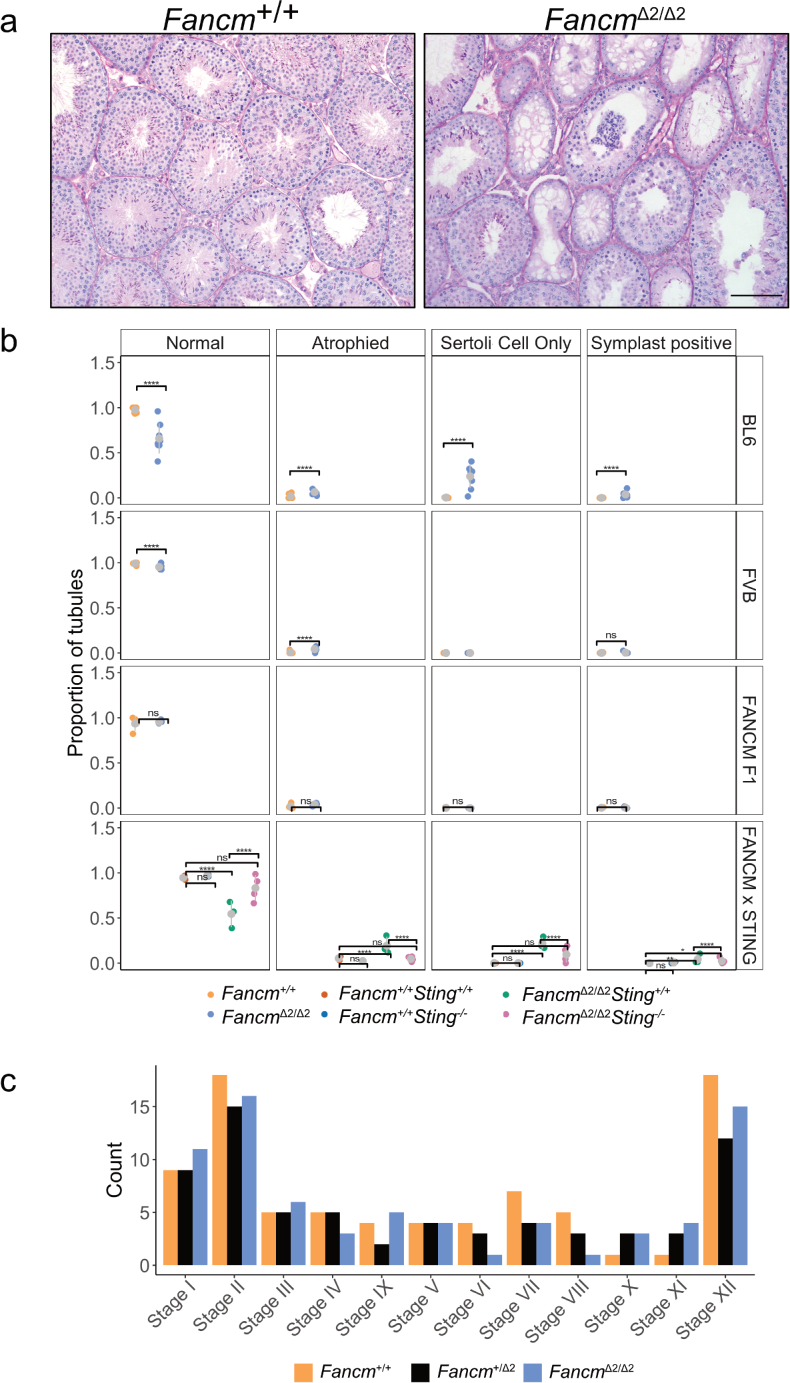
*Sting* depletion partially rescues histological defects in the seminiferous tubules in *Fancm* -deficient mice. a) PAS stain of seminiferous tubules in B6.*Fancm*^+/+^ and B6.*Fancm^Δ^*^2/^*^Δ^*^2^ mice. Scale bar = 100 *µ*m. b) Quantification of the PAS-labelled slides. Wild-type tubules are relatively replete in their appearance. Mutant mice have a heterogeneous reduction in germ cell content of the tubules. A mixture of normal, atrophied, symplast-positive and Sertoli cell-only tubules were be observed. c) Categorisation and quantification of seminiferous tubules of different spermatogenic stages in adult mice. **** indicates p-value *≤* 0.0001, ** indicates p-value *≤* 0.01, * indicates p-value *≤* 0.05 (pair-wised proportion test with multiple testing correction “fdr”). 127.625 *±* 19.87 B6.*Fancm^Δ^*^2/^*^Δ^*^2^ (8 mice), 174.13 *±* 15.62 B6.*Fancm*^+/+^ (8 mice), 109 *±* 31.39 FVB.*Fancm^Δ^*^2/^*^Δ^*^2^ (6 mice), 121 *±* 12.68 FVB.*Fancm*^+/+^ (6 mice), 168 F1.*Fancm^Δ^*^2/^*^Δ^*^2^ (4 mice), 161 *±* 82.76 F1.*Fancm*^+/+^ (4 mice), 145 *±* 15.56 *Fancm*^+/+^*Sting* ^+/+^ (2 mice), 158.5 *±* 9.19 *Fancm*^+/+^*Sting* ^-/-^ (2 mice), 159.67 *±* 17.5 *Fancm^Δ^*^2/^*^Δ^*^2^*Sting* ^-/-^ (3 mice), 151.25 *±* 10.34 *Fancm^Δ^*^2/^*^Δ^*^2^*Sting* ^-/-^ (4 mice) tubules counted.

**S19 Fig.**
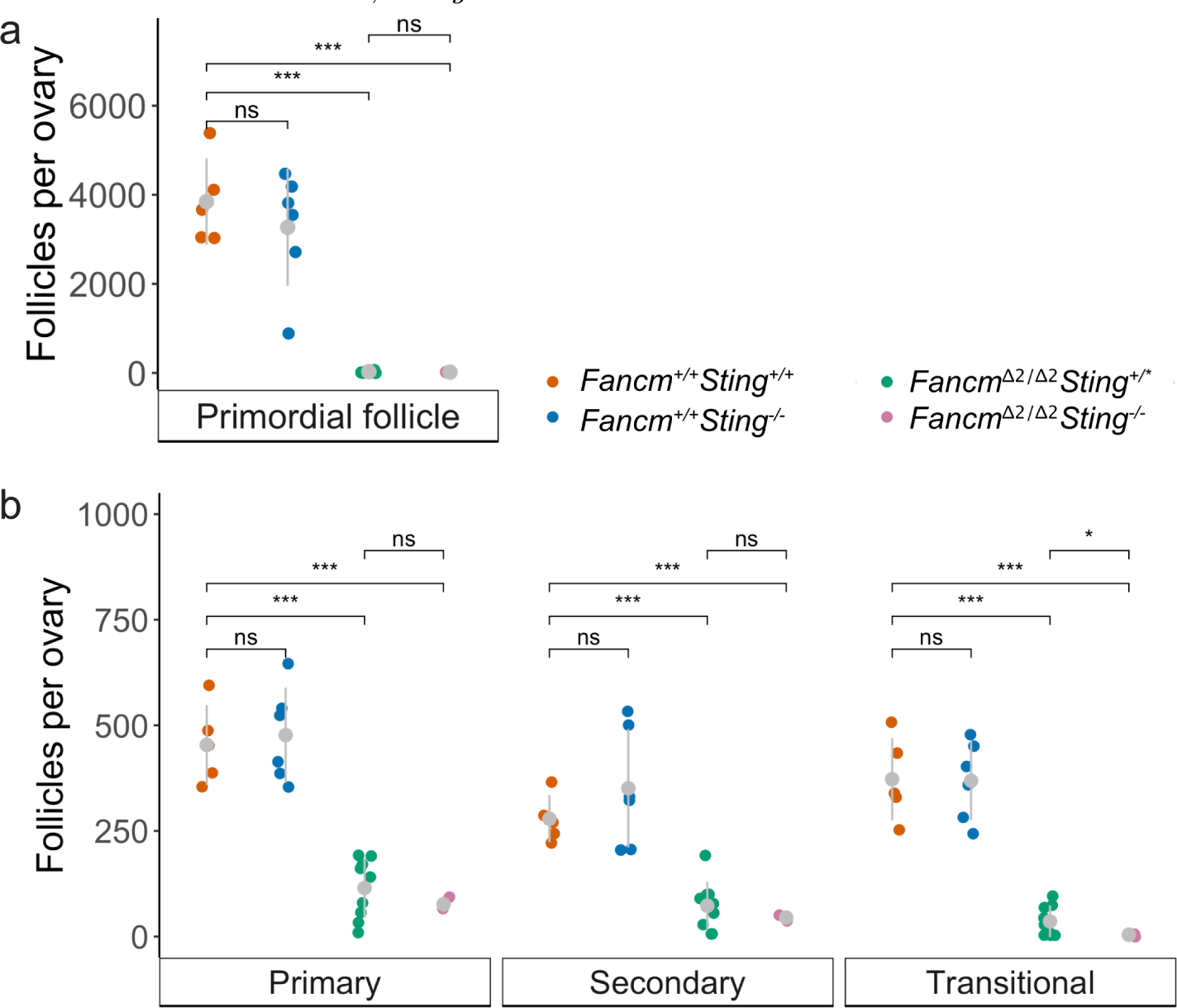
Sting depletion does not affect follicle numbers in *Fancm* deficient female mice. a) Primordial and b) growing follicles were assayed in mice deficient for *Fancm*, *Sting* or both.

**S20 Fig.**
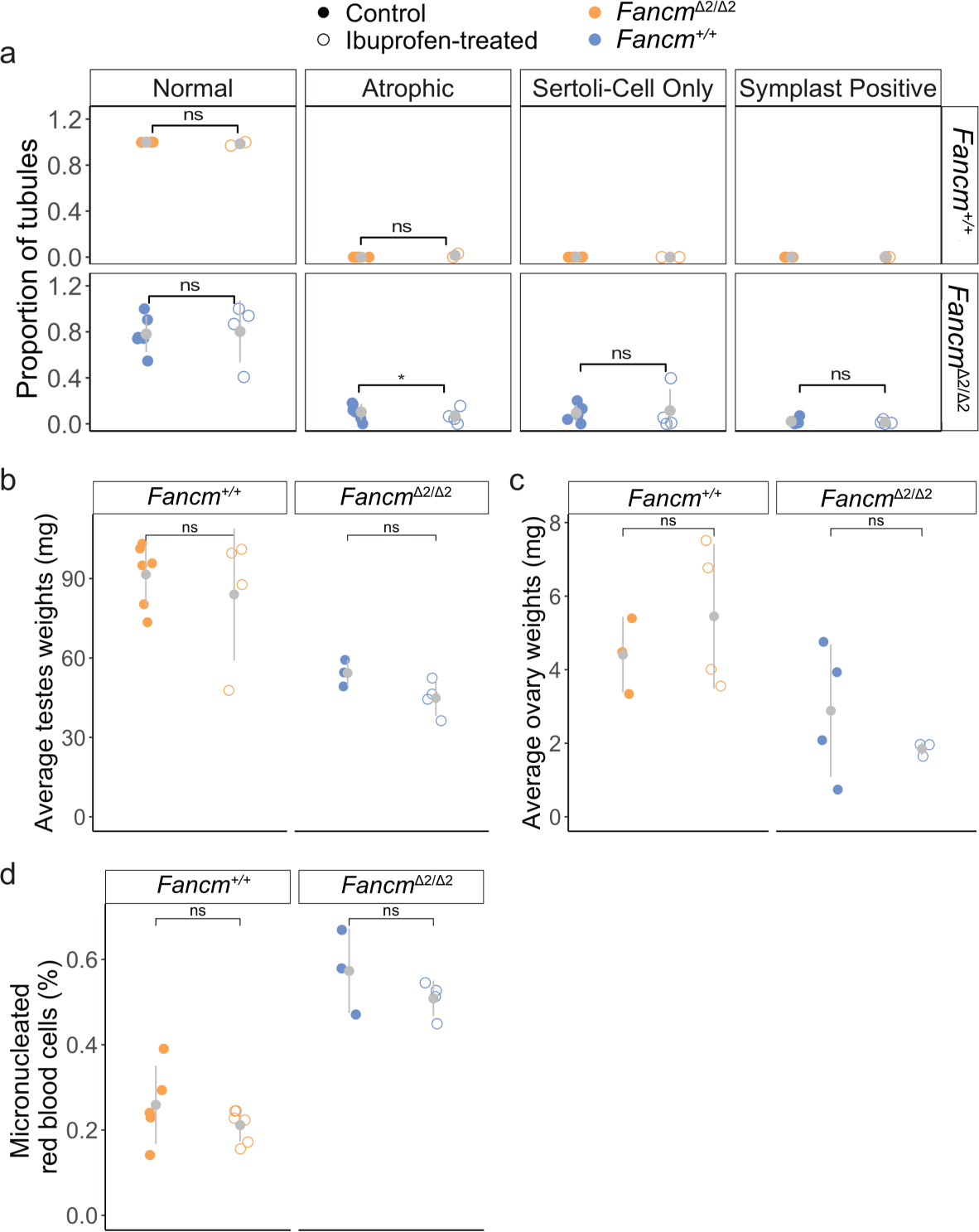
Ibuprofen does not improve gametogenesis, hypogonadism or genomic instability in *Fancm* -deficient mice. a) Quantification of H and E stain of seminiferous tubules in ibuprofen-treated B6.*Fancm^Δ^*^2/^*^Δ^*^2^ mice and controls b) Average testes weights of 8 week old ibuprofen-treated and untreated control mice c) Average ovary weights of 8 week old ibuprofen-treated and untreated control mice d) Genomic instability was assayed by quantification of micronucleated red blood cells of ibuprofen-treated and untreated control mice. **** indicates p-value *≤* 0.0001, ** indicates p-value *≤* 0.01, * indicates p-value *≤* 0.05 (pair-wised proportion test with multiple testing correction “fdr”). 16 *Fancm*^+/+^ and 24 *Fancm^Δ^*^2/^*^Δ^*^2^ control mice. 8 *Fancm*^+/+^ and 16 *Fancm^Δ^*^2/^*^Δ^*^2^ mice used.

**Table S1:** Offspring of FVB.*Fancm^Δ^*^2/+^ x FVB.*Fancm^Δ^*^2/+^ intercrosses.

|  | Male |  | Female |  |
| --- | --- | --- | --- | --- |
|  | Observed | Expected | Observed | Expected |
| *Fancm*<sup>+/+</sup> | 84 | 77 | 68 | 77 |
| *Fancm*<sup>Δ2/+</sup> | 161 | 153 | 149 | 153 |
| *Fancm*<sup>Δ2/Δ2</sup> | 70 | 77 | 81 | 77 |
| Total | 315 |  | 298 |  |

**Table S2:** Offspring of FVB.*Fancm^Δ^*^2/+^ x B6.*Fancm^Δ^*^2/+^ intercrosses.

|  | Male |  | Female |  |
| --- | --- | --- | --- | --- |
|  | Observed | Expected | Observed | Expected |
| *Fancm*<sup>+/+</sup> | 58 | 53 | 57 | 53 |
| *Fancm*<sup>Δ2/+</sup> | 115 | 106 | 92 | 106 |
| *Fancm*<sup>Δ2/Δ2</sup> | 56 | 53 | 45 | 53 |
| Total | 229 |  | 194 |  |

**Table S3:** Meiotic prophase substaging with SYCP3 is unchanged in F1.*Fancm^Δ^*^2/^*^Δ^*^2^ compared to F1.*Fancm*^+/+^. Data is from two mice per genotype.

|  | leptotene | zygotene | pachytene | diplotene | diakinesis |
| --- | --- | --- | --- | --- | --- |
| *Fancm*<sup>+/+</sup> | 1 | 35 | 160 | 31 | 8 |
| *Fancm*<sup>Δ2/Δ2</sup> | 9 | 24 | 170 | 36 | 11 |

**Table S4:** Meiotic metaphase univalent counts is unchanged in F1.*Fancm^Δ^*^2/^*^Δ^*^2^ compared to F1.*Fancm*^+/+^. Data is from seven and eight mice for the wild type and mutant respectively.

|  | balanced metaphase I | two unbalanced univalents |
| --- | --- | --- |
| *Fancm*<sup>+/+</sup> | 184 | 7 |
| *Fancm*<sup>Δ2/Δ2</sup> | 273 | 2 |

**Table S5:** B6.*Fancm^Δ^*^2/^*^Δ^*^2^ have absent populations of pachytene cells, which is not detected in B6.*Fancm*^+/+^. Data is from four mice of each genotype. Xsquared = 19.314, df = 1, p-value = 1.109e-05

|  | pachytene present I | pachytene absent |
| --- | --- | --- |
| *Fancm*<sup>+/+</sup> | 106 | 0 |
| *Fancm*<sup>Δ2/Δ2</sup> | 99 | 22 |

## Notes

### Competing Interest Statement

The authors have declared no competing interest.

